# The dynamic equilibrium of nascent and parental MCMs safeguards replicating genomes

**DOI:** 10.1101/828954

**Authors:** Hana Sedlackova, Maj-Britt Rask, Rajat Gupta, Chunaram Choudhary, Kumar Somyajit, Jiri Lukas

## Abstract

The MCM2-7 (minichromosome maintenance) protein complex is a DNA unwinding motor required for the eukaryotic genome duplication^1^. Although a huge excess of MCM2-7 is loaded onto chromatin in G1 phase to form pre-replication complexes (pre-RCs), only 5-10 percent are converted into a productive CDC45-MCM-GINS (CMG) helicase in S phase – a perplexing phenomenon often referred to as the ‘MCM paradox’^2^. Remaining pre-RCs stay dormant but can be activated under replication stress (RS)^3^. Remarkably, even a mild reduction in MCM pool results in genome instability^4, 5^, underscoring the critical requirement for high-level MCM maintenance to safeguard genome integrity across generations of dividing cells. How this is achieved remains unknown. Here, we show that for daughter cells to sustain error-free DNA replication, their mothers build up a stable nuclear pool of MCMs both by recycling of chromatin-bound MCMs (referred to as parental pool) and synthesizing new MCMs (referred to as nascent pool). We find that MCMBP, a distant MCM paralog^6^, ensures the influx of nascent MCMs to the declining recycled pool, and thereby sustains critical levels of MCMs. MCMBP promotes nuclear translocation of nascent MCM3-7 (but not MCM2), which averts accelerated MCM proteolysis in the cytoplasm, and thereby fosters assembly of licensing-competent nascent MCM2-7 units. Consequently, lack of MCMBP leads to reduction of nascent MCM3-7 subunits in mother cells, which translates to poor MCM inheritance and grossly reduced pre-RCs formation in daughter cells. Unexpectedly, whereas the pre-RC paucity caused by MCMBP deficiency does not alter the overall bulk DNA synthesis, it escalates the speed and asymmetry of individual replisomes. This in turn increases endogenous replication stress and renders cells hypersensitive to replication perturbations. Thus, we propose that surplus of MCMs is required to safeguard replicating genomes by modulating physiological dynamics of fork progression through chromatin marked by licensed but inactive MCM2-7 complexes.

---

Eukaryotic cells possess an efficient mechanism to restrict MCM assembly as pre-RCs only once per cell cycle^7^. In G1-phase, nearly an entire pool of MCM2-7 units are loaded onto the chromatin as pre-RCs (Extended Data Fig. 1a). A fraction of licensed pre-RCs is converted to CMGs and gives rise to active replisomes, which are critical for the complete and stable duplication of the genome^3^. Throughout S-phase, pre-RCs progressively dissociate from DNA^8^, either due to passive replication of unused, dormant origins or during fork termination when two active CMG units meet each other^9^. Since chromatin-detached MCMs are prone to degradation^10^ and the pre-RC formation starts already in late-mitosis^11^, we hypothesized the existence of a mechanism, which safeguards sufficient MCM levels in mother cells such that their daughters become fully competent for pre-RC licensing right from the start of their life cycles.

**Fig. 1.**
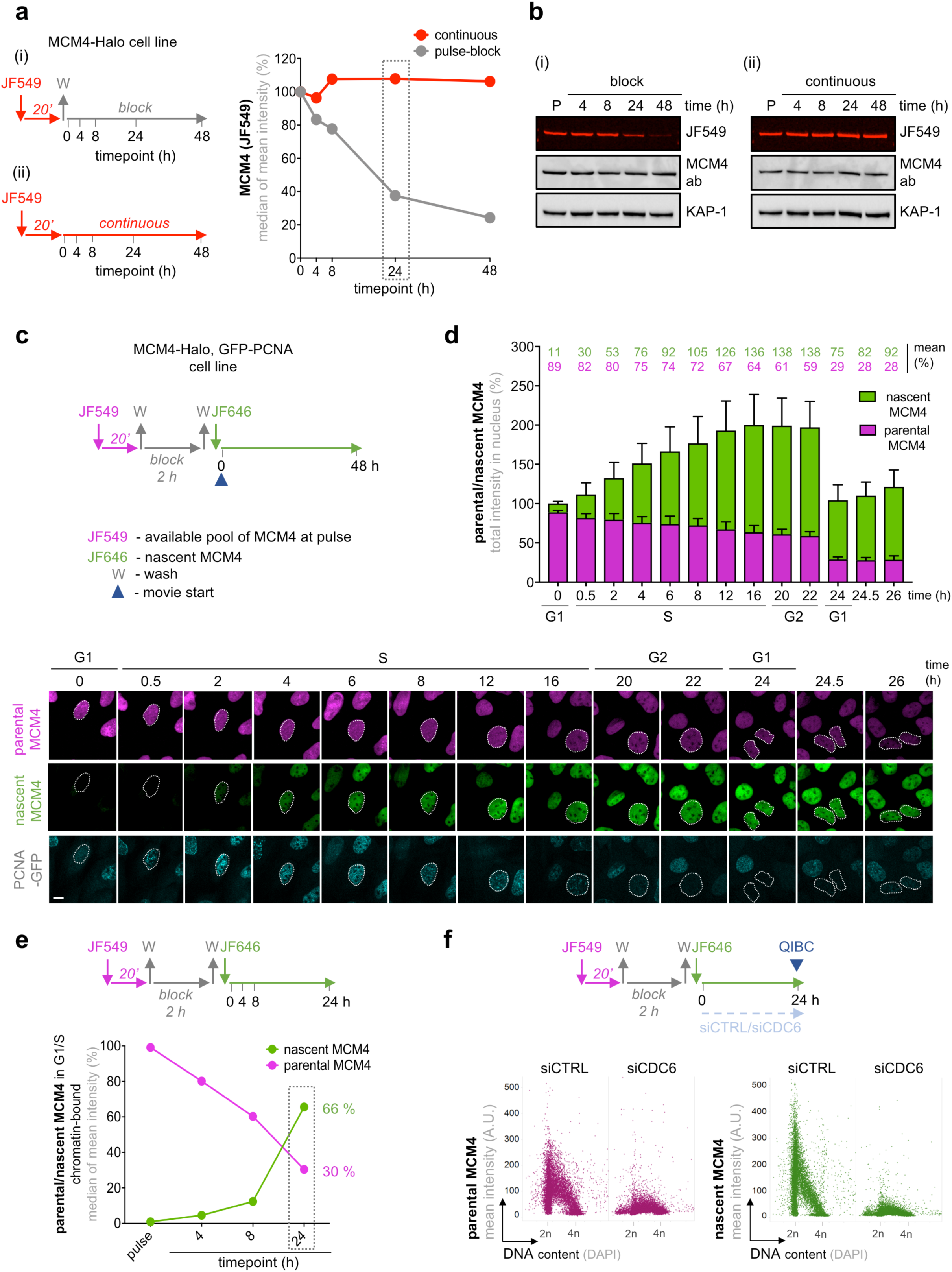
Continuous synthesis of nascent MCMs in mother cells mounts optimal origin licensing in daughter cells. **a,** left, MCM4 labeling protocol in U2OS cells endogenously expressing MCM4-Halo (i and ii). JF549, Janelia Fluor dye 549; W, wash. Right, QIBC-based quantification of MCM4-Halo at different intervals of Halo-Ligand labeling. n ≈3500 cells for each condition. See Extended data Fig. 1e. Box marks the cell doubling time. **b,** Western blotting (WB) of MCM4-Halo from an independent experiment performed as in (**a**). P: pulse. **c,** Top, MCM4-Halo dual labeling protocol. JF646, Janelia Fluor dye 646. Bottom, single plane projection (SPP) images of U2OS cells endogenously expressing MCM4-Halo and ectopically expressing GFP-PCNA with a dual labeling of MCM4-Halo. Scale bar, 14 μm. Dotted circles show a representative trajectory of parental (magenta) or nascent (green) MCM4 in an individual cell at indicated timepoints for one complete cell cycle marked by PCNA. Also see methods. **d,** Quantification of total intensity of MCM4-Halo fluorescence derived from the data in (**c**). Total intensity of parental and nascent of MCM4 at the start of time-lapse microscopy was pooled as 100 percent and represented as relative percentage. Each data point indicates mean ±SD. n = 15 cells. **e,** Top, MCM4-Halo dual labeling protocol. Bottom, quantification of G1/S-specific chromatin bound nascent and parental MCM4 levels. See also Extended data Fig. 3a. Box marks the cell doubling time. **f,** Top, MCM4-Halo dual labeling protocol. Bottom, QIBC of cells transfected with control or CDC6 siRNAs and stained for chromatin bound MCM4-Halo. Nuclear DNA was counterstained by 4′,6-diamidino-2-phenylindole (DAPI). Parental MCM4-Halo (left panel) and nascent MCM4 (right panel). n ≈10 000 cells for each condition. A.U., arbitrary units.

To test this hypothesis, we generated a HaloTag-MCM4 fusion construct and expressed it from the endogenous locus in human U2OS cells (Extended Data Fig. 1b). This enabled us to monitor MCM dynamics and stability in a defined genetic system and without adverse effects of protein overexpression. Using quantitative image-based cytometry (QIBC)^12^ of large cell populations, we first confirmed that a short pulse of fluorescent HaloTag ligand rapidly labeled the bulk of nuclear MCM4 including the fraction involved in licensed pre-RCs (Extended Data Fig. 1c, d). We then set out to investigate whether the MCM steady state levels are maintained by a dynamic fluctuation of the protein supply, shielding the old available pool, or combination of both. To mark the instantly available pool, we pulse-labeled MCM4 with fluorescent HaloTag ligand (JF549) for 20 minutes followed by releasing cells in a fresh media with a nonfluorescent HaloTag blocker to halt MCM labeling (Fig. 1a, left; i). Alternatively, to follow contribution of newly synthesized MCM4, we cultured the cells with continuous presence of the fluorescent HaloTag ligand for two rounds of cell division (48 hours) (Fig. 1a, left; ii). Strikingly, the nuclear levels of Halo-MCM4 gradually declined when chased with the blocker but remained stable during continuous labeling (Fig. 1a, right; Extended Data Fig. 1e). This was recapitulated by an immunoblot analysis of a fluorescent signal from HaloTag ligand (Fig. 1b). The total levels of MCM4 remained constant in either condition when analyzed by MCM4-specific antibody (Fig. 1b), indicating that the rapidly-declining pool of pulse-labeled MCM4 must have been replenished by a newly synthesized protein.

To test this prediction, we set out to directly visualize and quantify in real time the different pools of MCMs. We designed a dual-labeling protocol of MCM4 with two fluorescently labeled HaloTag ligands (JF549 and JF646, respectively) temporally separated with the HaloTag blocker in a U2OS cell line stably expressing green fluorescent protein (GFP)-tagged PCNA, a robust indicator of cell cycle progression^13^ (Fig. 1c). With this experimental setup, we could extend our previous findings by showing that MCM4 pulse-labeled with the JF549 ligand (which marks the pre-existing pool) steadily declined throughout the cell cycle (Fig. 1c, d). Furthermore, a chase with the JF646 ligand revealed a vivid production of new MCM4, starting from the S-phase entry and continuing until late-S/G2 (Fig. 1c, d). Intriguingly, at the end of the cell cycle, combination of the pre-existing and the newly synthesized protein doubled the total pool of MCM, ensuring that the newly-born daughter cells instantly receive the same total amount of MCMs, with which their mother started the previous cell cycle (Fig. 1d). Immunoblotting and QIBC-based analysis of Halo-MCM4 confirmed at large cell population levels a gradual loss of JF549 and progressive increase in JF646 pools before cell cycle completion (Extended Data Fig. 2a, b). Reassuringly, inhibiting protein synthesis by cycloheximide (CHX) or blocking the proteasome by MG132 confirmed that the stable nuclear pool of MCM is a result of progressive synthesis of JF646-MCM4, compensating for a gradual decay in JF549-MCM4 (Extended Data Fig. 2c, d). Reflecting these distinct features of the MCM pools, we name the JF549-labeled pool as ‘parental MCMs’, which have been at least once on chromatin and are recycled from the one cell cycle to the next. Following the same logic, we name the JF646-labeled pool as ‘nascent MCMs’, which were synthesized *de novo* by mother cells during S phase and were passed on to their daughters without previous engagement in pre-RCs. Together, our data suggest that despite their gradual decline, a fraction of parental MCMs is recycled for the next cell cycle. In parallel, the loss of the parental MCM pool is counterbalanced by native MCM synthesized throughout S phase of the maternal cell cycle.

To investigate the function of parental and nascent MCM pools inherited by daughter cells, we analyzed their respective contribution to chromatin-bound pre-RCs. QIBC analysis of chromatin-bound proteins revealed that the nascent and parental MCM4 were licensed in ∼2:1 ratio (Fig. 1e, Extended Data Fig. 3a, b), each via a canonical CDC6-dependent mechanism (Fig. 1f). Strikingly, although originating from different pools, both nascent and parental MCMs efficiently interacted with CDC45, suggesting an equal proficiency in forming an active CMG helicase in next cell cycle (Extended Data Fig. 3b). Very similar results were obtained with endogenous HaloTag-MCM2 subunit, indicating that efficient origin licensing in daughter cells critically relies on parental and nascent MCMs from the previous cell cycle (Extended Data Fig. 3d, e, f). These findings are aligned with a previous report in *S. cerevisiae* suggesting that the *de novo* generation of nascent Cdc6 and Mcm proteins during G1-phase drives each cycle of pre-RC formation^14^. Strikingly, however, unlike in budding yeast, we find that the human nascent MCMs are generated in the preceding S-phase in a fully licensing-competent mode, while being kept away from chromatin (and thus inducing re-replication) by degradation of MCM loader CDT1. Supporting this notion, treatment of cells with MLN4924, which stabilizes CDT1 by inhibiting cullin-RING E3 ubiquitin ligases^15^, resulted in re-licensing of both the parental and nascent MCMs in the same cell cycle (Extended Data Fig. 4a-c). Based on these data, we conclude that mother cells constantly replenish the gradual loss of parental MCMs by synthesizing nascent MCM subunits and thus ensure that daughter cells inherit sufficient amount of MCMs to sustain replication of their genomes. Consistent with this notion, QIBC analysis of endogenously tagged GFP-MCM2 and GFP-MCM4 confirmed a continuous increase in the total intensity of GFP signal (which unlike the mean intensity is independent of nuclear size/volume) as cells progress from G1 to G2 phase (Extended Data Fig. 4d).

Chromatin-bound MCMs are remarkably stable but much less is known about their turnover before they engage into licensed pre-RCs. Although MCM proteins do not require extensive folding by canonical chaperones such as HSP70/HSP90 (mainly attributed to their intrinsically globular secondary structure)^16^, they might need co-chaperones to rapidly reach their subcellular localization and assemble into stable MCM2-7 complexes. Intriguingly, MCM4 was shown to associate with a co-chaperone FKBP51 in a complex with MCMBP^16^, a hitherto poorly characterized ultrahigh affinity MCM interactor^6, 17^. We thus asked whether molecular chaperoning activity might assist to sustain the production of nascent MCMs complexes as wells as parental MCM complex released from chromatin in a given S phase. We focused on MCMBP, which is distantly related to MCMs and exhibits structure-function properties that are well suited to regulate MCMs throughout their life cycle^17^. We first generated U2OS cells ectopically expressing FLAG-MCMBP and analyzed the MCMBP interactome from the whole-cell extract by SILAC-based mass spectrometry (MS) (Extended Data Fig. 5a). Consistent with previous reports^6, 17, 18^, we found MCM subunits as top interactors of MCMBP, but we also noticed that MCMBP does not associate with the components of active CMG such as CDC45 or GINS4 (Extended Data Fig. 5a). We validated the MS data by reciprocal coimmunoprecipitation and immunoblot analysis (Extended Data Fig. 5b). Extended interaction analysis under more stringent ionic treatment of biochemically fractionated cell lysates (Extended Data Fig. 5c) revealed that MCMBP interacts with all the subunits of MCMs except MCM2 as described previosuly^6^, and is also refrained of active CMG components (Fig. 2a, b; Extended Data Fig. 5d). Furthermore, we noticed that the interaction between MCM and MCMBP was much more prominent in soluble fractions compared to chromatin-bound proteins (Fig. 2a, b), indicating that MCMBP might regulate a specific MCM pool that is distinct from pre-RCs or active CMG. To test this hypothesis, we used the CRISPR–Cas9 technology to generate a derivative of Halo-MCM4 U2OS cell line with endogenously-tagged GFP-MCMBP susceptible to an inducible auxin-based degradation (Extended Data Fig. 5e). This allowed us to analyze in an isogenic cellular system the fate of nascent and parental MCM4 after a rapid and quantitative MCMBP depletion (Fig. 2c, d). Strikingly, single-cell tracking revealed that while the dynamics of parental Halo-MCM4 remained unaltered, nascent Halo-MCM4 showed a massive delay in nuclear accumulation after MCMBP degradation (Fig. 2d, e). These observations were validated by QIBC (Extended Data Fig. 6a) and recapitulated in MCMBP knockout (MCMBP-KO) U2OS cell line (Extended Data Fig. 6b, c).

**Fig. 2.**
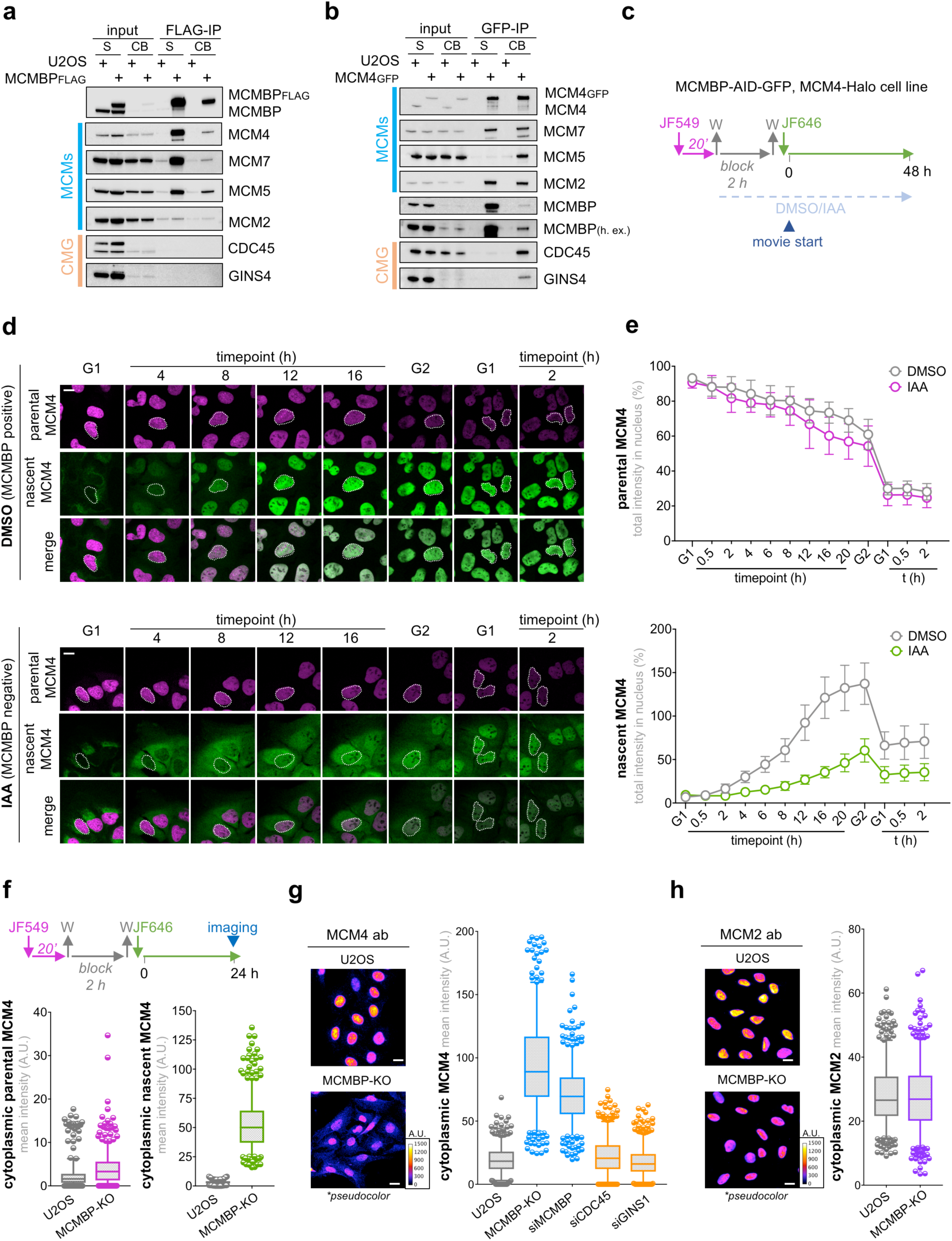
MCMBP stabilizes and translocates nascent MCM3-7 to cell nuclei. **a,** FLAG**-**immunoprecipitation (FLAG-IP) followed by western blotting of sub-cellular fractions from U2OS cells or its derivative stably expressing FLAG-tagged MCMBP S: supernatant, CB: chromatin bound. **b,** GFP-immunoprecipitation (GFP-IP) followed by western blotting of sub-cellular fractions from U2OS cells or its derivative endogenously expressing GFP-tagged MCM4. **c,** MCM4-Halo dual labeling protocol in U2OS cells endogenously expressing MCM4-Halo and MCMBP-GFP-degron. **d,** Representative SPP images of U2OS cells endogenously expressing MCM4-Halo and MCMBP-GFP-degron in the presence of DMSO (top) or auxin (IAA; bottom) with a dual labeling of parental (magenta)/nascent (green) MCM4-Halo in an individual cell (dotted circle) at indicated timepoints. Scale bar, 14 μm. See methods for information regarding cell cycle classification (G1 and G2). **e,** Quantification of total intensity of MCM4-Halo derived from the data in (**d**). Total intensity of parental (top) and nascent (bottom) of MCM4 at the start of time-lapse microscopy was pooled as 100 percent and represented as relative percentage for cells treated by DMSO or IAA. Each data point indicates mean ±SD. n = 15 cells. **f,** Top, dual-HaloTag labeling protocol in U2OS (MCM4-Halo) and MCMBP-KO (MCM4-Halo) cells. Blue triangle represents collection of cells for QIBC analysis. Bottom, mean fluorescence intensity (MFI) of cytoplasmic parental MCM4 (left) and nascent MCM4 (right) for indicated cells. The center line of the plot represents the median. The boxes indicate the 25th and 75th centiles, and the whiskers indicate 5 and 95 percent values. n = 500 per condition. **g,** Left, representative images of immunostained MCM4 in naïve U2OS and MCMBP-KO cells without pre-extraction. The color gradient indicates the mean MCM4 intensity. Scale bar, 20 μm. Right, quantification of MFI of cytoplasmic MCM4. n = 500 per condition. **h,** Left, images of immunostained MCM2 in naïve U2OS and MCMBP-KO cells without pre-extraction. The color gradient indicates the mean MCM2 intensity. Scale bar, 20 μm. Right, Quantification of MFI of cytoplasmic MCM2. n = 500 per condition.

Intriguingly, during the real-time tracking of MCMBP-deficient cells, we noticed that the paucity of nascent Halo-MCM4 in cell nuclei was accompanied by its accumulation in the cytoplasm (Fig. 2d), suggesting a possible role of MCMBP in nucleo-cytoplasmic trafficking of MCM proteins. Indeed, QIBC analysis of cells expressing Halo-labeled MCMs confirmed that the cytoplasmic mislocalization of MCM4 in MCMBP-KO cells was restricted to the nascent, but not parental MCM pool (Fig. 2f). Intriguingly, detailed analysis with antibodies specific to individual MCM subunits revealed that the lack of MCMBP severely compromised nuclear import of all MCMs except MCM2 (Fig. 2g, h; Extended Data Fig. 7a). Furthermore, siRNA-mediated depletion of MCMBP, but not CDC45 and GINS1, also recapitulated the cytoplasmic mislocalization of MCMs both in MCMBP-KO and MCMBP-degron cells (Fig. 2g), reinforcing a specific role of MCMBP in the nuclear trafficking of nascent MCMs.

From all MCM subunits, only MCM2 and MCM3 are known to possess an autonomous nuclear localization signal (NLS)^19^, leading to the assumption that MCM3-7 and MCM2 are transported across the nuclear membrane as two different units, similar to what was also reported for ORC sub-complexes^20^ or by a possible formation of multiple sub-complexes as observed upon overexpression of MCM2^19^. Moreover, a careful analysis of MCM3-7 nuclear translocation in MCMBP-negative cells suggests that while these MCM subunits were grossly mis-localized to the cytoplasm, a fraction of them eventually entered cell nuclei, albeit with slower kinetics (Fig. 2d). This led us to postulate that the MCM3-embedded NLS might not be sufficient to confer stable nuclear localization of the MCM3-7 subcomplex and that MCMBP might be required to boost nuclear import and retention. Indeed, bioinformatic analysis revealed a putative bipartite NLS motif in the N-terminus of MCMBP (Fig. 3a), whose deletion severely abrogated the nuclear import of MCMBP (Fig. 3a; Extended Data Fig. 7b) but also other MCM subunits (Fig. 3b; Extended Data Fig. 7c, d). Importantly, MCM2 nuclear localization remained independent of MCMBP nuclear transport (Fig. 3b). In support of this notion, MCMBP does not associate with MCM2 with high affinity when compared to the other MCM subunits^6^ (Fig. 2a).Together, these results establish that MCMBP translocates nascent MCM3-7 to the cell nuclei independent of the MCM2 subunit. Our findings are also in agreement with the observation that fission yeast Mcb1 (an ortholog of human MCMBP) plays a critical role in maintaining the nuclear localization of MCMs^17^. However, in contrast to the reported mechanism of Mcb1 in prohibiting aberrant nuclear export of MCM subunits^17^, our results elucidate the direct involvement of MCMBP in the nuclear trafficking of newly synthesized MCM3-7 subunits.

**Fig. 3.**
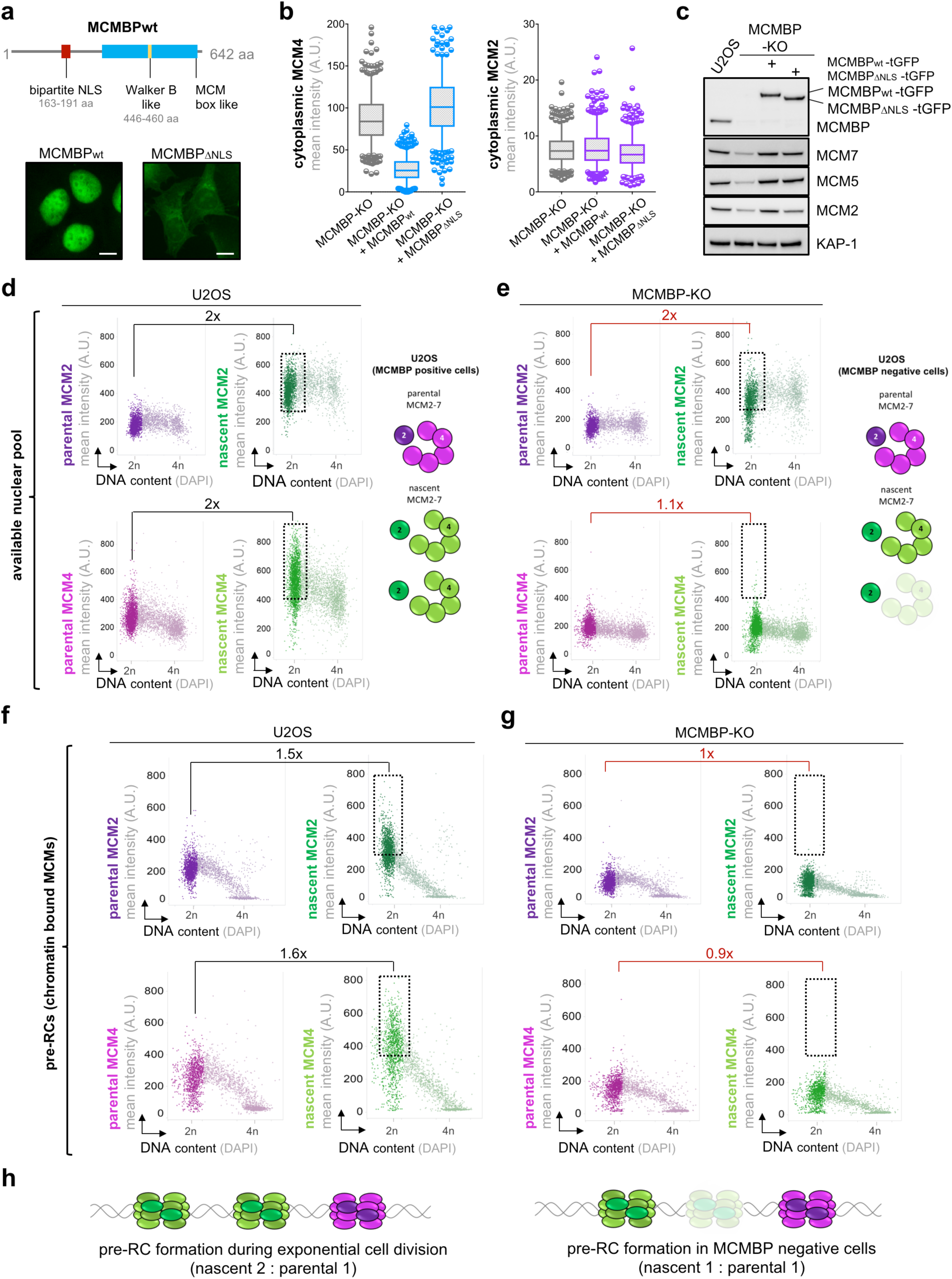
Daughter cells license pre-RCs by distinct pools of parental and nascent MCM subunits. **a,** Top, human MCMBP protein domains. Bottom, representative images of stably integrated GFP-tagged wt- and NLS-deleted MCMBP in MCMBP-KO cells. Scale bar, 14 μm. **b,** MFI of cytoplasmic MCM4 (right) and MCM2 (left). n = 500 per condition. The center line of the plot represents the median. The boxes indicate the 25th and 75th centiles, and the whiskers indicate 5 and 95 percent values. **c,** Western blotting of total cell extracts from naïve U2OS or MCMBP-KO cells complemented with either wt or NLS-deleted MCMBP. tGFP, turbo-GFP. **d,** Left, QIBC of naïve U2OS cells stained for parental (purple) or nascent (green) MCM2-Halo and DAPI (top), and parental (magenta) or nascent (green) MCM4-Halo and DAPI (bottom) without pre-extraction. Right, graphical summary of the inherited parental and nascent MCM pools in the naïve daughter cells. **e,** Left, QIBC of MCMBP-KO cells processed and analyzed as in (**d**). Right, graphical summary of the inherited parental and nascent MCM pools in MCMBP-KO daughter cells. Boxes in (**d**) and (**e**) mark the excess nascent MCM2 or MCM4 over the levels of parental MCM2 or MCM4 in G1/S phase. Data derived from Extended data Figs. 8b and 9b. **f,** Left, QIBC of naïve U2OS cells stained for chromatin bound parental (purple) or nascent (green) MCM2-Halo and DAPI (top) and chromatin bound parental (magenta) or nascent (green) MCM4-Halo and DAPI (bottom). **g,** Left, QIBC of MCMBP-KO cells processed and analyzed as in (**f**). Boxes in (**f**) and (**g**) mark the excess nascent MCM2 or MCM4 over the levels of parental MCM2 or MCM4 at G1/S phase. Data derived from Extended data Figs. 8c and 9c. **h,** Schematic outcome of pre-RC formation in naïve U2OS cells (left) and MCMBP-KO (right) with regard to distinct MCM2-7 complexes composed of nascent and parental subunits, respectively.

Next, we wanted to understand the fate of mislocalized cytoplasmic MCMs. Immunoblot analysis of MCM subunits in MCMBP-KO cells showed a marked reduction in total MCM pool, but not other replication-associated factors such as CDC45, GINS, TIMELESS, and PCNA (Extended Data Fig. 6b). The low levels of cytoplasmic MCM3-7 proteins were not associated with reduced transcripts (Extended Data Fig. 7e), indicating accelerated proteolysis. In support of this notion, treatment of MCMBP-deficient cells with the proteasome inhibitor MG132 stabilized the cytoplasmic pool of nascent MCM4 (Extended Data Fig. 7f). Strikingly, although localized in cytoplasm, the pool of MCMs remained stable in cells expressing the NLS-deficient MCMBP mutant as opposed to MCMBP-KO cells (Fig. 3c), indicating that a physical interaction of MCMBP with MCM3-7 is sufficient to shield the latter against proteolysis regardless of subcellular location. To our surprise, while MCM2 is transported to cell nuclei independently of MCMBP, its stability was compromised in the NLS-deficient MCMBP mutant (Fig. 3c), which is aligned with the alleviated nuclear as well as total protein levels of MCM2 in MCMBP-KO cells (Extended Data Fig. 6b, 7d). This indicates that under the conditions of reduced nascent MCM3-7 subunits, the unused MCM2 is also degraded, albeit with a slower kinetics (Extended Data Fig. 7g).

To further explore the relationship between distinct MCMs pools and their involvement in pre-RC formation, we set out to systematically analyze nascent and parental MCMs directly upon their inheritance by wild type and MCMBP-deficient daughter cells, respectively. To this end, we compared the total inherited nuclear pool of parental and nascent MCMs to their chromatin-bound fractions at the G1-S boundary, when the pre-RC licensing reaches its maximum levels. QIBC analysis of the total nuclear MCMs in daughter cells revealed a two-fold higher accumulation of nascent MCM2 as compared to parental MCM2, both in normal and MCMBP-deficient cells (Fig. 3d, e; Extended Data Fig. 8a, b). A very similar trend was observed also for MCM4 (used here as a proxy for MCM3-7 subcomplex) but only in MCMBP-proficient settings (Fig. 3d; Extended Data Fig. 9a, b). In sharp contrast, MCMBP-KO cells were unable to maintain this 2:1 ratio and instead featured almost equal levels of nascent and parental MCM4 subunits, creating a relative excess of nascent MCM2 over the other MCM subunits (Fig. 3e, Extended Data Fig. 9a, b). Strikingly, QIBC analysis of chromatin-bound fractions in MCMBP-KO daughter cells also revealed that the excess nascent MCM2 was not translated to increased origin licensing, suggesting that the superfluous nascent MCM2 was refrained from new pre-RC formation due to shortage of complementary pool of nascent MCM3-7 (Fig. 3g, Extended Data Fig. 8c, 9c). Notably, this result also indicated that nascent and parental MCMs might not mix in the same MCM2-7 units. This notion is based on a prediction that if nascent and parental MCMs readily mix, then the overabundant nascent MCM2 in MCMBP-KO cells would be expected to compete for its place both in nascent and parental MCM2-7 holocomplexes. However, QIBC analysis suggests the opposite by revealing a failure of nascent MCM2 to load on chromatin in MCMBP-deficient settings (Fig. 3g; compare rectangular boxes in Fig. 3e and 3g). Instead, as our previous data already indicated, the surplus level MCM2 excluded from chromatin under these conditions was gradually degraded (Fig. 2j, Extended Data Fig. 6b). Consequently, while in normal daughter cells, pre-RC licensing is composed of two copies of nascent and one copy of parental MCM2-7 rings, MCMBP negative daughter cells license only one copy of each, nascent and parental MCM2-7, resulting into sparsely organized pre-RCs (Fig. 3h). From these observations, MCMBP emerges as a multi-functional chaperone that confers stability to nascent MCM3-7 subunits and fosters their relocation to cell nuclei, and thereby supports the formation of licensing-competent MCM2-7 and thus contribute to reach the critical levels of pre-RCs.

Finally, to test the significance of nascent MCM production and maintenance in mother cells for genome integrity of the ensuing cell generations, we monitored total chromatin-bound MCMs as a readout for the efficiency of pre-RC formation. We consistently observed a dramatic loss of licensed pre-RCs in the absence of MCMBP (Fig. 4a). Surprisingly, in spite of the low level of chromatin-bound MCMs, the bulk DNA synthesis and chromatin association of active replisome components (e.g. TIMELESS and PCNA) remained very similar (Fig. 4b, c). To reconcile these opposing effects of MCMBP loss on pre-RCs and active replisomes, we monitored inter origin distance (IOD) to quantitively access origin density. Consistent with the reduced pre-RCs, we observed an increase in IOD in exponentially growing MCMBP-KO cells (Fig. 4d). Strikingly, when CLASPIN was depleted to boost the frequency of origin firing^21^, the enforced decrease in IOD was still evident in MCMBP-KO cells (Fig. 4d), suggesting that with such a low level of pre-RCs, MCMBP deficient cells still maintained a pool of dormant replication origins, although with a compromised density of initiation events. In line with these findings, we observed an increased frequency of 53BP1-nuclear bodies (Fig. 4e), an established hallmark of inheritable under-replicated DNA lesions arising at genomic loci lacking high density of replication origins^22, 23^.

**Fig. 4.**
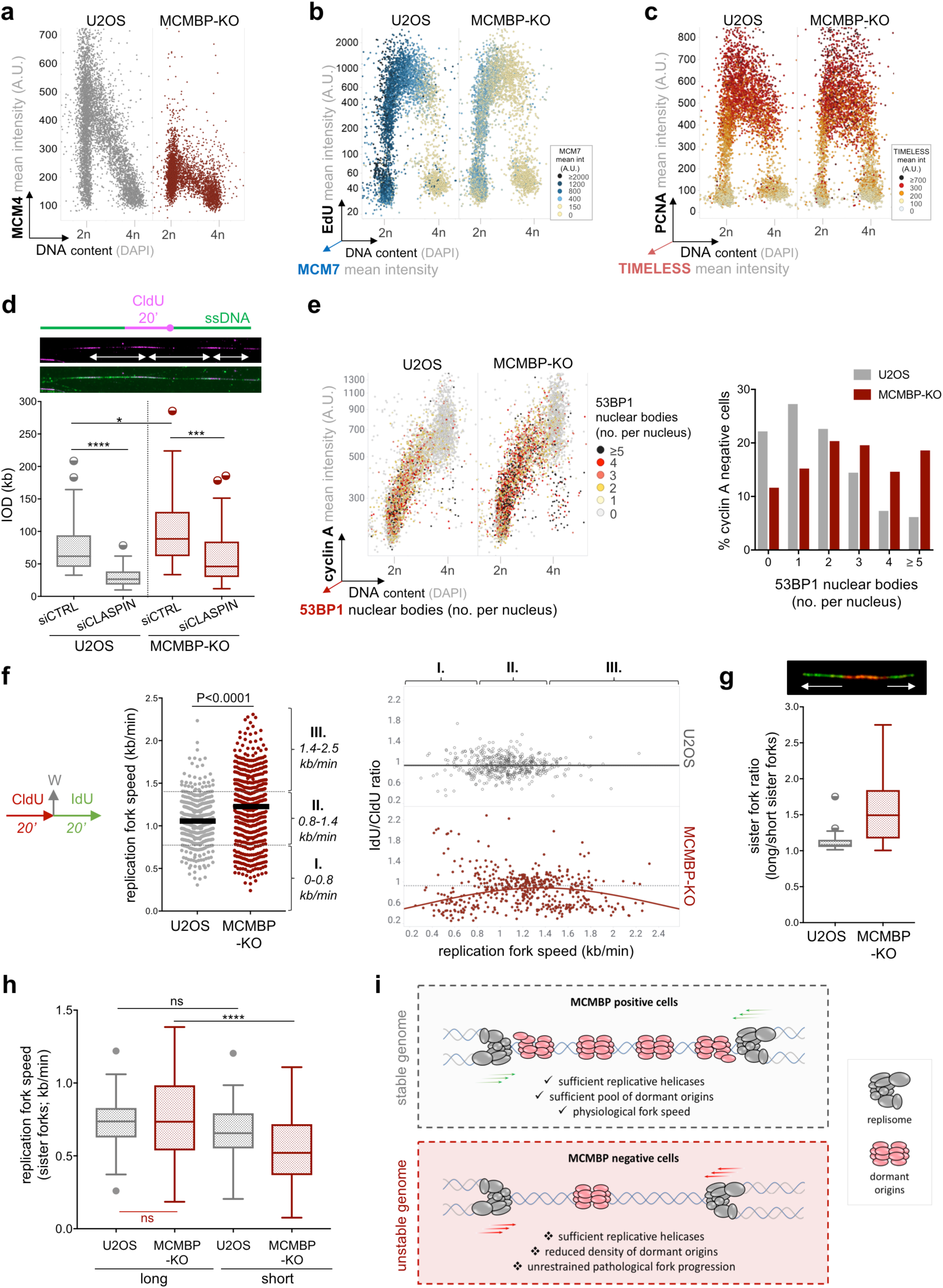
MCMBP loss restrains pre-RCs formation and induces replication stress. **a,** QIBC of MCM4 chromatin loading in U2OS or MCMBP-KO cells as indicated. n ≈3500 cells per condition. **b,** QIBC of EdU incorporation in naïve or MCMBP-depleted U2OS stained for MCM7 and DAPI. n ≈5000 cells for each condition. The color gradient indicates the mean intensity of chromatin-loaded MCM7. **c,** QIBC of TIMELESS in U2OS or MCMBP-KO cells co-stained for PCNA and DAPI. n ≈3500 cells for each condition. The color gradient indicates the mean intensity of chromatin-bound TIMELESS. **d,** Top, DNA fiber labeling protocol to monitor inter-origin distance (IOD). ssDNA, single stranded DNA. Bottom, IOD in U2OS or MCMBP-KO cells treated with control or CLASPIN siRNAs. The central line of the box and whisker depict the median of Tukey plot. The boxes indicate the 25th and 75th percentiles. n = 60 initiation events. (*, P=0.0259; ***, P=0.0007; ****, P< 0.0001; ns, not significant) **e,** Left, QIBC of 53BP1 nuclear bodies (NBs) in U2OS or MCMBP-KO cells co-stained for cyclin A and DAPI to discriminate cell cycle phases (n ≈5700 cells for each condition; colors indicate the number of 53BP1 nuclear bodies per nucleus). Right, Quantification of 53BP1 NBs in the depicted cell populations. **f,** Left, DNA fiber labeling protocol. Middle, replication fork speed in cells as indicated. The line represents median. n = 500 fibers. Right, individual fork ratio is derived from the data in (left) by dividing the length of DNA tracts labeled by IdU and CldU, respectively. The grey (U2OS) and red (MCMBP-KO cells) lines represent Gaussian fitting. **g,** Top, a representative example of asymmetrical bi-directional fork. Bottom, quantification of sister fork ratio from 50 bidirectional forks for each condition. **h,** Fork speed derived from the long and short sisters of bidirectional replication forks (from data in Fig 4g). (****, P< 0.0001; ns, not significant) **i,** A model depicting the critical role of MCM surplus to support optimal levels of replicative helicases, dormant origins, and physiological fork speed across multiple ensuing cell divisions.

Next, to understand whether the reduced pre-RCs directly impact DNA replication at the individual fork level, we measured fork speed using the DNA fiber technique^24^. Strikingly, we found that the absence of MCMBP (and the corresponding reduction of pre-RC licensing) mildly increased the overall rate of replication fork progression (Fig. 4f). Acceleration of replication forks under the reduced levels of chromatin-bound MCMs was further supported by partial depletion of the MCM loader CDT1 (Extended Data Fig. 10a), suggesting that abundance of origin licensing is tightly associated with the physiological progression of the replisome. Based on these results, we postulated that a surplus of chromatin-loaded inactive MCM2-7 complexes could provide physical resistance to the moving forks, absence of which might unleash uncontrolled fork progression and increase the frequency of pausing or stalling events. Of note, although median fork speed showed a shift towards faster forks, we consistently observed a prominent population of slow-moving forks in MCMBP-KO cells (Fig. 4f). Consistent with the idea that abnormal fork progression can impose stress on replicating genomes^25^, we found increased levels of individual fork asymmetry, both at the slow and the fast end of the fork speed spectrum in MCMBP-KO cells (Fig. 4f). Importantly, both fork acceleration and asymmetry in MCMBP-KO cells could be rescued by reintroducing wildtype, but not NLS-deficient MCMBP mutant (Extended Data Fig. 10b, c), suggesting that the paucity of chromatin-loaded MCMs directly impacts the physiological movement of individual forks and cause replication-associated stress. To further test this prediction, we monitored sister fork asymmetry, a direct readout for the replication stress arising due to uneven processivity on either side of the replication bubble^25^. Supporting this idea, MCMBP-KO cells showed increased incidence of asymmetry in bidirectional forks (Fig. 4g), suggesting frequent pausing of individual replisomes. To understand which pool of fork speed was responsible for causing asymmetrical extension of replication bubble in MCMBP deficient cells, we evaluated speed of sister forks (derived from the tract lengths on either side of bidirectional forks). Strikingly, while comparison of shorter sister tracts showed substantial slowing of replication forks in MCMBP specific manner, long sister tracts between normal and MCMBP-KO cells exhibited remarkably similar fork rates (Fig. 4h). This imbalance in speed of long and short tracts skewed the overall symmetry of bidirectional replication in MCMBP deficient cells (Fig. 4g). However, to our surprise, a complete absence of accelerated sister forks in MCMBP deficient cells (Fig. 4h; as opposed to unidirectional replication forks in Fig. 4f) directly implied that the unrestrained speed of replication forks led to their frequent pausing/stalling and turning them into shorter sister tracts, and also explains the incidence of slow forks observed in Fig. 4f (population I). Furthermore, stress at the individual fork level was accompanied by elevated levels of global chromatin-bound RPA, activation of ATR signaling, increased micronuclei formation, and massive sensitization to topoisomerase I inhibitor CPT, which are often associated with hallmarks of replication stress^26^ (Extended Data Fig. 10d-h). Together, these data suggest that MCMBP is required to sustain MCM levels at the threshold required for to maintain optimal origin density and physiological fork speed.

Collectively, this study uncovers how cells generate and maintain surplus of MCM subunits across ensuing generations to alleviate endogenous DNA replication stress. The salient new addition to the current understanding is our finding that MCM pool is sustained by continuous recycling of already licensed parental MCMs, and a simultaneous synthesis of the nascent pool already in mother cells, thereby ensuring that daughter cells receive sufficient amount of licensing-competent MCM units as soon as they enter the new cell cycle (Extended Data Fig. 10i). Perhaps most strikingly, while we find that both parental and nascent MCMs retain equal proficiency for pre-RC formation and subsequent activation as functional CMG helicases, our results also indicate that nascent and parental MCM subunits do not readily mix to form ‘chimeric’ MCM complexes. The functional consequences of this are not clear at this point but these findings open up a new avenue to study whether parental MCMs involved as pre-RCs in the previous cell cycle are inherited by daughter cells with specific post-translational modifications that pre-determine their biochemical activity. Conceptually, these findings are broadly analogous to old and new H3-H4 dimers that remain in distinct pools upon nucleosome disruption and reassembly during DNA replication^27^. Mechanistically, we uncovered a specific requirement of MCMBP in safeguarding the contribution of nascent MCM3-7 subunit to overall MCM pool in mother cells before they divide. Without MCMBP, daughter cells inherit only half of the nascent MCM2-7 units, which results into drastically impaired licensing of replication origins (Fig. 4i, Extended Data Fig. 10i). Interestingly, recent studies showed that cells released from quiescence enter the first S-phase with severely underlicensed chromatin and are thus particularly vulnerable to replication stress^28, 29^. Based on our findings, we suggest that this is partly caused the fact that every time a cell commits to proliferate, it needs to pass the first S phase to build sufficient amount of nascent MCMs and thus sustain the ensuing cell cycles with minimum endogenous replication stress. Strikingly, MCMBP mediated maintenance and licensing of excess MCMs is largely dispensable for exponential DNA synthesis as well as preservation of dormant ‘back-up’ origins. In this regard, our findings allowed us to revisit a long-standing enigma called the “MCM paradox”^30^ by postulating that beyond its role in supplying backup replication origins under stressed conditions, the high surplus of MCMs is vital to enforce physiological pace of replication fork progression (Fig. 4i, Extended Data Fig. 10i). Thus, we propose an unanticipated role of inactive chromatin-loaded MCM2-7 as an inbuilt genome surveillance mechanism to set the physiological threshold of fork speed and limit replication-associated stress (Fig. 4i). From this perspective, we define ‘fork-speed management’ as one on the main functions of 10-20 fold excess DNA-bound MCMs, a concept that can illuminate the notion that even a mild alteration in MCM2 or MCM4 levels are associated with the increased incidence of spontaneous tumor formation^31, 32^. Alterations in physiological fork progression and accumulation of spontaneous replication-associated stress might explain the extreme tumor susceptibly penetrance of hypomorphic variants of MCMs^31, 32^. Furthermore, based on our discovery of the critical role of MCMBP in nascent MCM maintenance, we propose that pharmacological inhibition of MCMBP may sensitize cancer cells by increasing their endogenous burden of replication stress due to pathologically accelerated forks, and decreased density of potential ‘back-up’ replication origins.

## Acknowledgements

Research funding was provided by the Novo Nordisk Foundation (grant NNF14CC0001), the Danish Cancer Society (grant R204 A12615) to J.L. H.S. was supported by the Novo Nordisk Foundation (grant NNF16CC0020906). K.S. was supported by the Danish Council for Independent Research (grant EDFF-FSS 82262) and the Lundbeck Foundation (grant R264-2017-2819). R.G. was supported by a European Molecular Biology Organization long-term postdoctoral fellowship (ALTF271-2014). C.C. was supported by the Hallas Møller Investigator Fellowship from the Novo Nordisk Foundation (NNF14OC0008541) and by the European Union’s Horizon 2020 research and innovation program (grant 648039). We thank J. Bulkescher and J. Dreier from the Protein Imaging Platform for their assistance with microscopy and image analysis. FACS analyses were carried out at CPR and Danstem Flow Cytometry Platform. The pX335 and pX458 plasmids were a gift from F. Zhang, MLN4924 was a gift from J. Duxin, and siRNAs against CDC45 and GINS1 were kindly gifted by L. Toledo. CHO cell line ectopically expressing MCM4-Emerald and RFP-PCNA was a gift from D. Gilbert. We sincerely thank C. Lukas for conceptual and technical inputs to this study and members of the Lukas lab for stimulating discussions and critical comments on the manuscript.

## Author contributions

H.S. K.S. and J.L. conceived the project and planned the study. H.S. performed experiments and prepared figures. H.S generated all the cell lines with the help of M.-B.R. K.S. performed IOD analysis. R.G. carried out proteomic data acquisition under the supervision of C.C. H.S. K.S. and J.L. analyzed the data and wrote the manuscript. All authors read and commented on the manuscript.

## Materials & correspondence

Should be addressed to K.S. or J.L.

## Methods

### Cell culture

The human U2OS osteosarcoma cell line (authenticated by STR profiling, IdentiCell molecular diagnostics) were grown in Dulbecco’s modified Eagle’s medium (DMEM, high glucose, Glutamax) containing 10% FBS and penicillin-streptomycin antibiotics (Thermo Fisher Scientific), under standard cell culture conditions (5% CO2, humidified atmosphere). All cell lines used and generated in this study were routinely tested for mycoplasma contamination (MycoAlert, Lonza).

### Cell lines

#### CRISPR/Cas9 generation of endogenously tagged cell lines

U2OS cells expressing C-terminally endogenously tagged proteins of interest were generated using CRISPR-Cas9 mediated homology-directed repair as described^33, 34^. Paired guide RNAs (gRNA) for specified genomic locus (MCM2: guide#1 TAGGGCCTCAGAACTGCTGC and guide#2 GCCATCCATAAGGATTCCTT, MCM4: guide#1 AAGGCTTCAGAGCAAGCGCA and guide#2 CTGCTTGCTGCACGCCACAT, CDC45: guide#1 GCATCAGGGTCGGGCTCTGA and guide#2 GCTCTGTCCTCCCTCAACGG) were inserted into pX335-U6-Chimeric_BB-CBh-hSpCas9n(D10A) (Addgene plasmid #42335, a gift from Feng Zhang) via *Bbs*I restriction site. For generation of MCMBP-AID-mEGFP cell line, single guide RNA (GTAATACCTATGAAGAGTAA) was cloned into pX458-pSpCas9(BB)-2A-GFP (gift from Feng Zhang, Addgene plasmid #48138) via *Bbs*I restriction site. U2OS cells were transfected by Lipofectamine LTX Plus reagent (Thermo Fisher Scientific, 15338-100) according to manufacturer’s recommendations with plasmids (pX335/pX458) containing cloned gRNA and donor plasmid containing the tag (mEGFP/AID-mEGFP/Halo) with flexible linker flanked by 900 bp homology arms complementary to the C-terminus of specific gene. Transfected cells were expanded before cell sorting of GFP-positive cells to obtain population of U2Os cells expressing proteins tagged by mEGFP. For generation of cells expressing Halo-Tag, cells were pulsed for 30 min with cell permeable TMR-552 ligand (Promega, G8251) in final labeling concentration (1 μM) followed by 30 min wash with fresh DMEM media before cell sorting. After 5 days, sorted cells were serially diluted into 100-mm dishes to obtain single isolated colonies. Individual colonies representing clonal cell population were isolated and expanded for their further characterization by both, western blotting (with antibody against GFP/Halo and MCM2/MCM4/CDC45/MCMBP), and junction PCR at specified genomic locus followed by Sanger sequencing. 2-3 clones of each cell line with homozygous tagging of all alleles were further functionally validated by immunofluorescence (sub-cellular localization of tagged-protein in a direct comparison with antibody-based staining) and immunoprecipitation (where interaction of tagged proteins and its key partners were tested). Only cell lines which passed all validation steps were used in final experiments.

#### MCMBP-KO cell line

Knock-out of MCMBP gene in U2OS cells was generated using single gRNA (AGGGGAACTTCGTTCAGTGA - targeting exon 3) or (AAATGGAGTTAATCCTGACT – targeting exon 2) cloned into pX458-pSpCas9(BB)-2A-GFP via *Bbs*I restriction site followed by Lipofectamine LTX Plus transfection. After 2 days, transfected cells were sorted for GFP-Cas9 positive cells. After 5 days, sorted cells were serially diluted into 100-mm dishes to obtain single isolated colonies. Clonal cell lines were expanded and further tested for knock-out of MCMBP gene by western blot (with antibody against MCMBP) and Sanger sequencing of gRNA targeting sites. Three cell clones for each gRNA containing knock-out of MCMBP gene were selected for phenotype testing. All tested clones showed the same phenotype and are represented by 2 clones (#1 and #2) in this study.

MCMBP-KO cell line expressing C-terminally Halo-tagged MCM2/MCM4 were generated with the same procedure as described above. For complementation assays, turboGFP-MCMBPwt (Origine NM_024834) or turboGFP-MCMBPΔNLS (bipartite NLS was identified using cNLS Mapper^35^ generated by site directed QuickChange II XL Site-Directed Mutagenesis kit (Agilent, 200522) with primers (forward: GTCCCTCAACATCCTACACTCCTAGTGGGAGTGTTGGTGGTCTTC and reverse: CCATTGAAGACCACCAACACTCCCACTAGGAGTGTAGGATGTTG) were transfected using Lipofectamine LTX Plus reagent into MCMBP-KO cells. Next day, transfected cells were serially diluted into 100-mm dishes and selected with DMEM medium containing 400 μg/ml Geneticin (Gibco, 10131-027) for approx. 12 days to obtain single isolated colonies. Individual colonies were isolated and transferred to 24-well plates. Clonal cell lines were expanded and further tested by fluorescence microscopy for MCMBP cellular localization and the level of expression was tested by western blot (with antibody against MCMBP/tGFP).

#### MCMBP-degron cell line

MCMBP-AID-mEGFP cell line for auxin induced MCMBP degradation (expressing C-terminally AID-mEGFP-tagged MCMBP) was generated and validated for homozygous tagging of all alleles with the procedure as described above. Afterward, cells were transfected using Lipofectamine LTX Plus reagent with plasmid (pCMV6-A-puro-TIR1-9xMyc) which contains codon-optimized (specific for human) TIR1 gene (paralog of Arabidopsis thaliana AFB2 gene). Next day, transfected cells were serially diluted into 100-mm dishes and selected with DMEM medium containing puromycin (1 μg/ml; Gibco, A11138-03) for 2-3 weeks to obtain single isolated colonies. Individual colonies were isolated and expanded. Ectopic expression of TIR1 was tested by immunofluorescence and western blot (using antibody against Myc). MCMBP degradation was achieved by the addition of indole-3-acetic acid (IAA; Sigma-Aldrich, I5148-10G) in final concentration 0.5 mM in fresh DMEM media. MCMBP-AID-mEGFP/TIR1-Myc cell line expressing C-terminally Halo-tagged of MCM4 was generated with the same procedure as described above.

#### Cell lines with ectopically expressing proteins

U2OS cells endogenously expressing C-terminal mEGFP-tagged MCM2 were transfected with plasmid (pCellCycleChromobody-RFP) containing RFP-PCNA chromobody (Chromotek, ccr) encoding single chain antibody recognizing endogenous PCNA protein. Single clones were selected under continuous growth in puromycin (1 μg/ml).

By the same procedure, stably overexpressing GFP-PCNA chromobody (pCellCycleChromobody-GFP) were introduced into U2OS cells containing endogenously Halo-tagged MCM4 and MCMBP-KO cells with endogenously Halo-tagged MCM4.

For generation of U2OS cells expressing FLAG-MCMBP, cells were transfected with FLAG3x-MCMBP subcloned from MCMBPwt-turboGFP. Single clones were selected with Geneticin (G418, 10131-027, 400 μg/ml).

### Drugs and Supplements

Cycloheximide (Sigma-Aldrich, C7698-1G, 12.5 μg/ml), MG132 (Calbiochem, 474790-10MG, 2 μM), campthotecin (Sigma-Aldrich, 208925-50MG), MLN4924 (R&Dsystems, I-502-01M, 5 μM) were used for indicated timepoints. CldU and IdU (Sigma-Aldrich) and EdU (Thermo Fisher Scientific, 31985070) were used as indicated.

### Gene silencing by siRNA

Cell transfection with siRNAs (Ambion Silencer Select) was performed using Lipofectamine RNAiMax (Thermo Fisher Scientific, 13778075) at a concentration of 10 nM MCMBP (s36586), CDC6 (s2744), CDC45 (s15829), GINS1 (custom made with sequence for #sense AAAACCAGUCUGAUGUGAAU[dT][dT] and #antisense AUUCACAUCAGACUGGUUUU [dT][dT]), 5 nM CLASPIN (s34330) and 1 nM CDT1 (s37723). Non-targeting siRNA (Ambion negative control #1) was used as control siRNA in all experiments.

### Antibodies

Antibody for immunofluorescence (IF) or western blot (WB) were used as follows: 53BP1 (mouse, Milipore, MAB3802, 1:1000 for IF), alpha-tubulin (mouse, Santa Cruz, sc-5286, 1:1000 for WB), CDC45 (rabbit, Cell Signaling Technology, 11881S, 1:1000 for WB), CDT1 (rabbit, Abcam, ab202067, 1:1000 for IF), cyclin A (rabbit, Santa Cruz, sc-751, 1:500 for IF), FLAG M2 (mouse, Sigma-Aldrich, F1804, 1:1000 for WB), GFP (rabbit, Chromotek, PABG1-100, 1:1000 for WB), turboGFP (rabbit, Thermo Fischer Scientific, PA5-22688, 1:1000 for WB), GINS4 (rabbit, Novus Biologicals, NBP2-16659, 1:1000 for WB), H2AX-phospho-S139 (mouse, Biolegend, 613401, 1:1000 for IF), H3 (rabbit, Abcam, ab1791, 1:5000 for WB), Halo (mouse, Promega, G9211, 1:1000 for WB), KAP-1 (rabbit, Bethyl Laboratories, A300-274A, 1:2000 for WB), MCM2 (mouse, Novus Biologicals, H000041171-M01, 1:1000 for IF, 1:1000 for WB), MCM3 (mouse, Santa Cruz, sc-390480, 1:500 for IF, 1:1000 for WB), MCM4 (mouse, Novus Biologicals, H00004173-B01P, 1:500 for IF, 1:500 for WB), MCM5 (rabbit, Abcam, ab17967, 1:1000 for IF, 1:2000 for WB), MCM6 (rabbit, Novus Biologicals, NBP1-82642, 1:200 for IF, 1:1000 for WB), MCM7 (mouse, Santa Cruz, sc-9966, 1:1000 for IF, 1:1000 for WB), MCMBP (rabbit, Novus Biologicals, NBP1-90746, 1:500 for IF, 1:2000 for WB), Myc (mouse, Abcam, ab32, 1:1000), PCNA (human, Immuno Concepts, 2037, 1:500 for IF), PCNA (mouse, Santa Cruz, sc-56, 1:1000 for WB), RPA32-phospho-S33 (rabbit, Bethyl Laboratories, A300-246A, 1:500 for IF), RPA70 (rabbit, Abcam, ab79398, 1:1000 for IF), TIMELESS (rabbit, Abcam, ab109512, 1:500 for IF, 1:1000 for WB).

Secondary antibody conjugates for IF were goat anti-mouse and goat anti rabbit Alexa Fluor 488 (A11029, A11034), Alexa Fluor 568 (A11031, A11036) Alexa Fluor 647 (A21236, A21245) (all from Thermo Fischer Scientific, 1:1000) and donkey anti-human Alexa Fluor 647 (Jackson Immuno Research, 709-605-149, 1:1000). Secondary antibody conjugates for WB were HRP horse anti-mouse IgG antibody (Vector Laboratories, PI-2000, 1:10000) and HRP goat anti-rabbit IgG antibody (Vector Laboratories, PI-1000, 1:10000).

### Western blot

To obtain whole cell extracts, cells were incubated with lysis buffer (10 mM HEPES pH 7.5, 500 mM NaCl, 1 mM EDTA, 1% NP-40) supplemented with protease and phosphatase inhibitors (ROCHE) and benzonase (Sigma-Aldrich, E1014-25KU) followed by analysis by NuPAGE 4-12% Bis-Tris gel (Thermo Fischer Scientific) after boiling samples in reducing buffer containing DTT as per standard procedures. Primary antibodies were diluted in PBS-Tween containing 5% powdered milk and incubated overnight at 4 °C. Secondary peroxidase-coupled antibodies were incubated for 1 hour at room temperature. ECL-based chemiluminiscence reagent (Amersham, RPN2106) was used for detection with an Odyssee-Fc system.

### Immunoprecipitation (IP)

For IP from whole cell extracts, U2OS cells expressing MCMBP-FLAG or MCM4-GFP or CDC45-GFP were harvested and lysed in RIPA buffer (Sigma-Aldrich, R0278-500ML) supplemented with protease and phosphatase inhibitors and benzonase. Whole cell extracts were incubated with anti-FLAG M2 magnetic beads (Sigma-Aldrich, F7425) or GFP-Trap magnetic beads (Chromotek, gtma-20) for 2 hrs at 4 °C. To elute bound proteins, beads were incubated with 200 μg/ml 3x FLAG peptides (Sigma-Aldrich, F4799-4MG) for 2 hrs at 4 °C. In case of GFP-trap, bound proteins were eluted by β-mercaptoethanol for 30 min at 95 °C. The immunoprecipitates were then analyzed with western blot with antibodies against indicated proteins or processed for mass spectrometry analysis. For IP from soluble and chromatin fraction, the subcellular fractionation was performed from U2OS expressing MCMBP-FLAG or MCM4-GFP or CDC45-GFP. The harvested cell pellets were resuspended in hypotonic buffer (10 mM HEPES pH 7.5, 1.5 mM MgCl_2_, 5 mM KCl, 1 mM DTT, 0.5% NP40) supplemented by protease and phosphatase inhibitors, EtBr, 5% glycerol and RNaseA and incubated 2 min on ice and then centrifuged at 16000 g for 5 min. The soluble fraction was collected and adjusted to 500 mM NaCl to maintain the same salt concentration as in chromatin fraction. Next, pellet was washed by washing buffer (10 mM HEPES pH 7.5, 5 mM NaCl, 0.3 M sucrose supplemented by protease and phosphatase inhibitors) and centrifuged at 16000 g for 5 min. The washing step was repeated twice. Finally, the pellets were resuspended in chromatin-lysis buffer (10 mM HEPES pH 7.5, 500 mM NaCl, 5 mM KCl, 1mM EDTA, 1% NP40) supplemented by protease and phosphatase inhibitors, EtBr, 5% glycerol, RNaseA and benzonase and incubated 45 min on ice followed by sonication at low amplitude and then centrifuged at 16000 g for 30 min. Soluble and chromatin fractions were applied on anti-FLAG M2 or GFP-Trap magnetic beads with the same procedure as described above.

### SILAC-based mass spectrometry and analysis of MCMBP-interactome

For SILAC experiments, naïve U2OS cells were grown in medium containing unlabeled L-arginine and L-lysine (Arg0/Lys0) as the light condition and U2OS cells expressing MCMBP-FLAG were grown in medium containing isotope-labeled variants of L-arginine and L-lysine (Arg10/Lys8) as the heavy condition. FLAG-IP was performed as described above. Proteins eluted from beads were boiled in 30 μl 4x NuPAGE LDS sample buffer (Invitrogen) containing 1 mM DTT followed by alkylation with 5.5 mM chloroacetamide. Next, the proteins were resolved on NuPAGE Novex Bis-Tis 4-12 % gel (Invitrogen), the gel was stained with Novex colloidal blue stain (Invitrogen) and subsequently destained with water. Lanes for each sample were sliced and destained further with a buffer containing 25 mM ammonium bicarbonate and 50% ethanol. Dehydration of gel pieces was done by addition of 100% ethanol followed by protein in-gel digestion with trypsin (Sigma-Aldrich) at 37 °C for 16 hrs. The gel pieces were treated with trifluoroacetic acid and the resulting peptides were eluted with increasing concentration of acetonitrile and desalted on reversed-phase C18 StageTips^36^. Peptides were eluted from StageTips by 40 μl of elution buffer containing 60% acetonitrile and 0.1% trifluoroacetic acid and then acetonitrile concentration was reduced in the eluates to less than 5 % by vacuum centrifugation. Before injecting into mass spectrometer, the peptides were diluted with buffer containing 0.5% acetic acid and 0.1% trifluoroacetic acid. The raw data files were analyzed using MaxQuant (version 1.5.2.8). Parent ion and MS/MS spectra were searched using Andromeda search engine^37^, a database against human proteome obtained from the UniProtKB (released in February 2012). To search for tandem mass spectra following settings were used: mass spectra tolerance of 6 ppm (MS mode), mass tolerance of 20 ppm (HCD MS2 mode), strict trypsin specificity and maximum 2 missed cleavages were allowed. N-terminal protein acetylation, and methionine oxidation were searched as variable modifications, whereas cysteine carbamidomethylation was searched as a fixed modification. The dataset was filtered based on posterior error probability (PEP) to arrive at a false discovery rate of below 1% estimated from a target-decoy approach. Table with SILAC ratio were then exported and analyzed in TIBCO Software to generate rank plot for MCMBP-interactome.

### HaloTag ligands and labeling protocol

For HaloTag labeling protocol (i) used in Fig. 1a, b and Extended Data Fig. 1e, cells expressing MCM4-Halo were pulsed with Janelia Fluor 549 (JF549) HaloTag ligand (Promega, GA1111) in final labeling concentration 200 nM for 20 min, washed three times with fresh DMEM medium and incubated fresh DMEM medium containing non-fluorescent blocking ligand in final labeling concentration 100 μM for indicated timepoints. Non-fluorescent blocking ligand was prepared as described^38^. Briefly, 100 mM HaloTag Succinimidyl Ester (O4) ligand was incubated with 500 mM Tris-HCl (pH 8.0) for 60 min at 25 °C to mask the functional groups. For HaloTag labeling protocol (ii) used in Fig. 1a, b and Extended Data Fig. 1e, cells expressing MCM4-Halo were incubated with JF549 HaloTag ligand in final labeling concentration 200 nM for indicated timepoints.

For dual-HaloTag labeling protocol, U2OS/MCMBP-degron/MCMBP-KO cells expressing MCM4-Halo/MCM2-Halo were incubated with JF549 HaloTag ligand in final labeling concentration 200 nM for 20 min, washed three times with fresh DMEM medium and incubated DMEM medium containing non-fluorescent blocking ligand in final labeling concentration 100 μM for 2 hours. After incubation, non-fluorescent blocking ligand was washed out and cells were additionally washed three times with fresh DMEM medium and incubated DMEM medium containing Janelia Fluor 646 (JF646) HaloTag ligand (Promega, GA1121) in final labeling concentration 200 nM for indicated timepoints.

### Immunofluorescence (IF) staining

Cells were grown on round 12 mm diameter, 1.5 mm thick glass coverslips (cleaned in 96 % ethanol, dried and autoclaved; Menzel-Glaser, 6307356). For immunostaining of chromatin bound proteins, cells were pre-extracted with ice-cold PBS containing 0.2% TritonX-100 for 2 min on ice before fixation 4% buffered formaldehyde for 15 min at room temperature. For immunostaining of nuclear pool of proteins, cells were without pre-extraction fixed with 4% buffered formaldehyde for 15 min at room temperature. When HaloTag labeling protocol was performed, cells were incubated with indicated HaloTag ligands for specified timepoints (for details see HaloTag ligands and labeling protocol, and schematic protocols in figures) before fixation (with/without preceding pre-extraction). When Click-iT EdU staining was performed, cells were incubated with 10 μM EdU for 20 min before pre-extraction and fixation. EdU detection was performed according to the manufacturer’s recommendations (Thermo Fisher Scientific) before incubation with primary antibodies. Primary and secondary antibodies were diluted in DMEM medium containing 10% FBS and 0.05% sodium azide (filtered through a 0.2 μm filter) and incubated at room temperature for 90 min and 30 min, respectively. For DAPI staining, secondary antibody solution was supplemented with 4’,6’-diamidino-2-phenylindole-dyhydrochloride (DAPI, 0.5 μg/ml). After staining, coverslips were washed three times with PBS and additionally twice in distilled water, dried and mounted with Mowiol-based mounting medium (Mowiol 488 (Calbiochem), glycerol, Tris-HCl pH 8.5).

### Quantitative image-based cytometry (QIBC)

QIBC was performed as previously described^12, 39^. Images were acquired using ScanR inverted high-content screening microscope (Olympus) equipped with wide-field optics, UPLSAPO dry objective (20x, 0.75-NA), fast excitation and emission filter-wheel devices for DAPI, FITC, Cy3 and Cy5 wavelengths, an MT20 illumination system and a digital monochrome Hamamatsu ORCA-R2 CCD camera (yielding a spatial resolution of 320 nm per pixel at 20x and binning of 1). Images were acquired in an automated fashion with the ScanR acquisition software (Olympus 2.7.1). At least 2000 cells per condition were acquired. Acquired images were processed and analyzed with ScanR analysis software. A dynamic background correction was applied to all images. The DAPI signal was used for the generation of an intensity-threshold-based mask to identify individual nuclei as main objects. This mask was then applied to analyze pixel intensities in different channels for each individual nucleus. After segmentation of nuclei, 53BP1-NB were segmented as above, and the desired parameters for the different nuclei or foci were quantified, with single parameters (mean and total intensities, foci count, and foci intensities) as well as calculated parameters (sum of foci intensity per nucleus). Table with values was then exported and analyzed in TIBCO Software to quantify absolute, median and average values in cell populations and to generate color-coded scatter plots. Within one experiment, similar cell numbers were compared for the different conditions and for visualization jittering was applied (random displacement of objects along the x axis) to make overlapping markers visible. The mean fluorescence intensity of cytoplasmic MCMs was quantified with ImageJ software.

### Confocal 3D imaging of live cells

Time-lapse imaging was acquired using an UltraVIEW Vox spinning-disk microscope (Perkin Elmer) and Volocity software (v.6.3) with a 40x, 1.3-NA Plan-Apochromat oil immersion objective and appropriate excitation and emission filters. Images were captured using a Hamamatsu EMCCD 16-bit camera at a sampling resolution of 121 nm in the x, y dimensions and 250 nm in the z dimension. Laser power and exposure time were appropriately adjusted with identical settings applied within series of experiments. Microscope performance and channel alignment were regularly checked via the imaging of 200 nm multicolor fluorescent beads.

For live cell time-lapse imaging, cells were seeded at appropriate density in imaging dishes (Nunc, Lab-Tek, 155361) and dual HaloTag labeling protocol (see details above) was performed up to end of incubation with non-fluorescent blocking ligand. Next after washing steps, the JF646 ligand diluted in CO_2_-independent medium was added to the cells followed by overlaid with mineral oil to minimize evaporation. Time-lapse imaging was acquired under stable temperature conditions of 37 °C, at 12 different positions using autofocusing (Nikon Eclipse TI microscope equipped with Nikon Perfect Focus System) and z-stacks (300 nm distance, 15 slices). Recording was performed for 48 hours with 30 min intervals. Laser power, exposure times and acquisition intervals were chosen appropriately to minimize sample bleaching. All images displayed in figures represent single plane projection (SPP) from the center of a 3D stack. Brightness and contrast were linearly adjusted for optimal presentation for each condition.

Nascent and parental MCMs were monitored from G1 phase to the next G1 phase using PCNA to differentiate between individual phases. G1 phase was determined by homogenous smooth nuclear distribution of PCNA intensities with heterogenous pattern for parental MCMs reflecting their loading on chromatin. S phase was determined by the onset and cessation of clearly discernible PCNA foci and G2 phase with homogenous smooth nuclear distribution of PCNA and parental MCMs reflecting their eviction from chromatin during S phase. Total intensities of nascent and parental MCMs were measured using ImageJ software from G1 phase (1^st^ timepoint) to the next G1 phase. After background correction, total intensities of nascent and parental MCM4 at 1^st^ timepoint were sum up and taken as 100 percent for calculation of relative percentage of nascent and parental MCMs in following timepoints separately for U2OS cells or MCMBP-KO cells (or in case of DMSO or IAA treatment in MCMBP degron cells). For the analysis of MCM dynamics in MCMBP degron cells (with DMSO or IAA treatment), G1 phase was defined based on heterogenous MCM pattern reflecting their loading on chromatin (parental MCM4), and G2 phase was demarcated based on the reverse time points (3-4 frames of time-lapse imaging) preceding mitosis.

### Confocal microscopy

Confocal imaging was carried out on an LSM 880 microscope with Airyscan (Zeiss AxioObserver. Z1) equipped with an oil immersion objective alpha Plan-Apochromat 100x/1.3 DIC M27. Images were acquired in super-resolution mode using Airyscan detector with appropriate emission filters for each laser line. Images were processed with deconvolution algorithm in LSM-ZEN software.

### Fluorescence recovery after photobleaching (FRAP)

For FRAP, U2OS cells expressing MCM2-mEGFP and RFP-PCNA were seeded at appropriate density in imaging dishes (Nunc, Lab-Tek, 155361) and before imaging DMEM medium was changed for CO_2_-independent medium. RFP-PCNA used to differentiate between individual phases of cell cycle. FRAP was acquired using an UltraVIEW Vox spinning-disk microscope (Perkin Elmer) with a 60x, 1.4-NA Plan-Apochromat oil immersion objective under stable temperature conditions of 37 °C. Volocity software (v.6.3) was used for FRAP setup. After 10 pre-bleaching frames (pre), a single bleach pulse (488-nm argon laser set to 100% power) was delivered in defined region (approximately 5 μm in diameter) followed by time-lapse for 35 seconds at maximum imaging scan (6 frames per second) with the laser transmission attenuated to 2.5%. Subsequently, the mean GFP-associated fluorescence intensity was extracted for each timepoint in the following regions: bleaching region (I_frap_(t)), background fluorescence outside the nucleus (I_back_(t)) and fluorescence intensity within the nucleus in which bleaching was performed (I_ref_(t))^40^. After background correction, double normalization (equation 1) which corrects for differences in the starting intensity in I_frap_ region and for loss in total nuclear fluorescence in I_ref_ region due to the bleaching pulse and to acquisition bleaching.

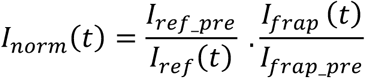

where

I_norm_(t) - normalized intensity

I_frap_pre_ / I_ref_pre_ - average of mean intensity in the I_frap_ / I_ref_ regions before bleach moment

Next, full scale normalization (equation 2) was applied to corrects for differences of the bleaching efficiencies (all recovery curves start from 0).

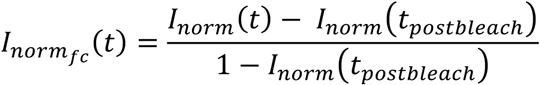

where

I_normfc_(t) - full scale normalized intensity

I_norm_(t_postbleach_) - is the first post-bleach value of the double normalized data

Mean of full scale normalized FRAP intensities were plotted from 14 cells per phase of cell cycle.

### RT-PCR

Total RNA from U2OS, MCMBP depleted cells using siRNA and MCMBP-KO (clone #1 and #2) cells was isolated using RNeasy Mini Kit (Qiagen, 74104). The cDNA was synthesized using High-Capacity cDNA Reverse Transcription Kit (Thermo Fischer Scientific, 4368814) followed by real-time PCR using primers (MCM2 forward: TGCAAGCCAGGAGACGAGA reverse: CCATTGGCAGTGTTGAGGG, MCM5 forward: ATTGGCTCCCAGGTGTCTGA reverse: GCGAGTCCATGAGTCCAGTG, MCM7 forward: CCCCTCTTTCTCCCATGCTG reverse: AGGCCCAGGCTAGAAGATGA) and Brilliant II SYBR Green qPCR Master Mix (Agilent, 600828).

### DNA fibers

DNA fibers were performed under same procedure as previously described^25^. Antibodies for DNA fibers were used as follows: for the tracts labeled with CldU (anti-BrdU, rat, Abcam, ab6326, 1:100) and IdU (anti-BrdU mouse, Becton Dickinson, 347580, 1:200). Secondary antibodies were goat anti-rat Alexa Fluor 594 IgG (Thermo Fischer Scientific, A21209, 1:100) and goat anti-mouse Alexa Fluor 488 IgG (Thermo Fischer Scientific, A11029, 1:100).

For IOD measurements, labeled cells were diluted 1/10 in non-labeled ones prior to fiber preparation. For anti-ssDNA (Tecan/IBL International 18731, 1:500) antibody was used.

### Clonogenic survival assay

Clonogenic survival experiments were performed as previously described^25^.

### Statistical analysis

All statistical analysis was done using unpaired student t-test or one-way ANOVA in GraphPad Prism v.7.0b.

---

**Extended Data Fig. 1.**
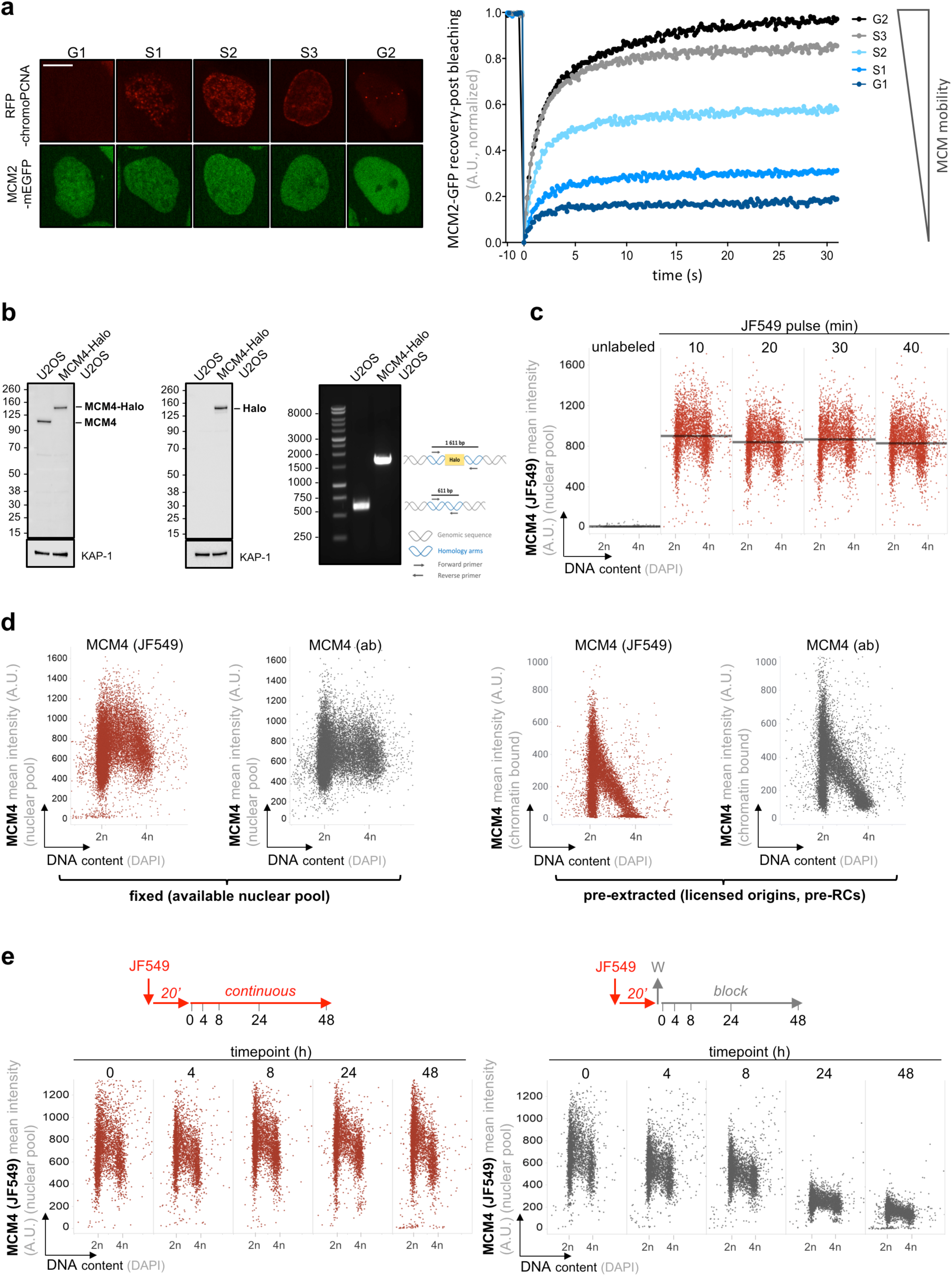
Development of tools and characterization of endogenous MCM proteins. **a,** Left, representative images of U2OS cells with endogenously GFP-tagged MCM2 and ectopically expressing RFP-chromoPCNA at indicated cell cycle stages used for FRAP analysis. Scale bar, 14 μm. Right, a summary of the MCM2-GFP FRAP curves at indicated cell cycle stages. n = 14 per cell cycle stage. **b,** Left, U2OS whole-cell lysates with endogenously tagged MCM4-Halo immunoblotted with MCM4 antibody. Middle, U2OS whole-cell lysates with endogenously tagged MCM4-Halo immunoblotted with Halo antibody. KAP-1 was used as loading control. Right, junction PCR showing homozygous MCM4-Halo tagging. **c,** QIBC of MCM4-Halo cells pulsed with JF549 HaloTag ligand (200 nM) for indicated time points. Nuclear DNA was counterstained by 4′,6-diamidino-2-phenylindole (DAPI). The line represents median. n ≈3500 cells per condition. A.U., arbitrary units. **d,** Left, QIBC of MCM4-Halo cells pulsed with JF549 HaloTag ligand (200 nM) for 20 min and immunostained for MCM4 without pre-extraction before fixation. Nuclear DNA was counterstained by DAPI. n ≈16000 cells per condition. Right, QIBC of MCM4-Halo cells pulsed with JF549 HaloTag ligand (200 nM) for 20 min and immunostained for MCM4 and DAPI with pre-extraction before fixation. n ≈12000 cells per condition. **e,** Left, HaloTag labeling protocol in MCM4-Halo cells. QIBC quantification of MCM4-Halo cells continuously labeled with 200 nM JF549 HaloTag ligand at indicated timepoints. Right, HaloTag labeling protocol in MCM4-Halo cells. QIBC-based quantification of MCM4-Halo cells pulsed with 200 nM JF549 HaloTag ligand followed by addition of 100 μM non-fluorescent blocking ligand for indicated timepoints. n ≈3500 cells per condition.

**Extended Data Fig. 2.**
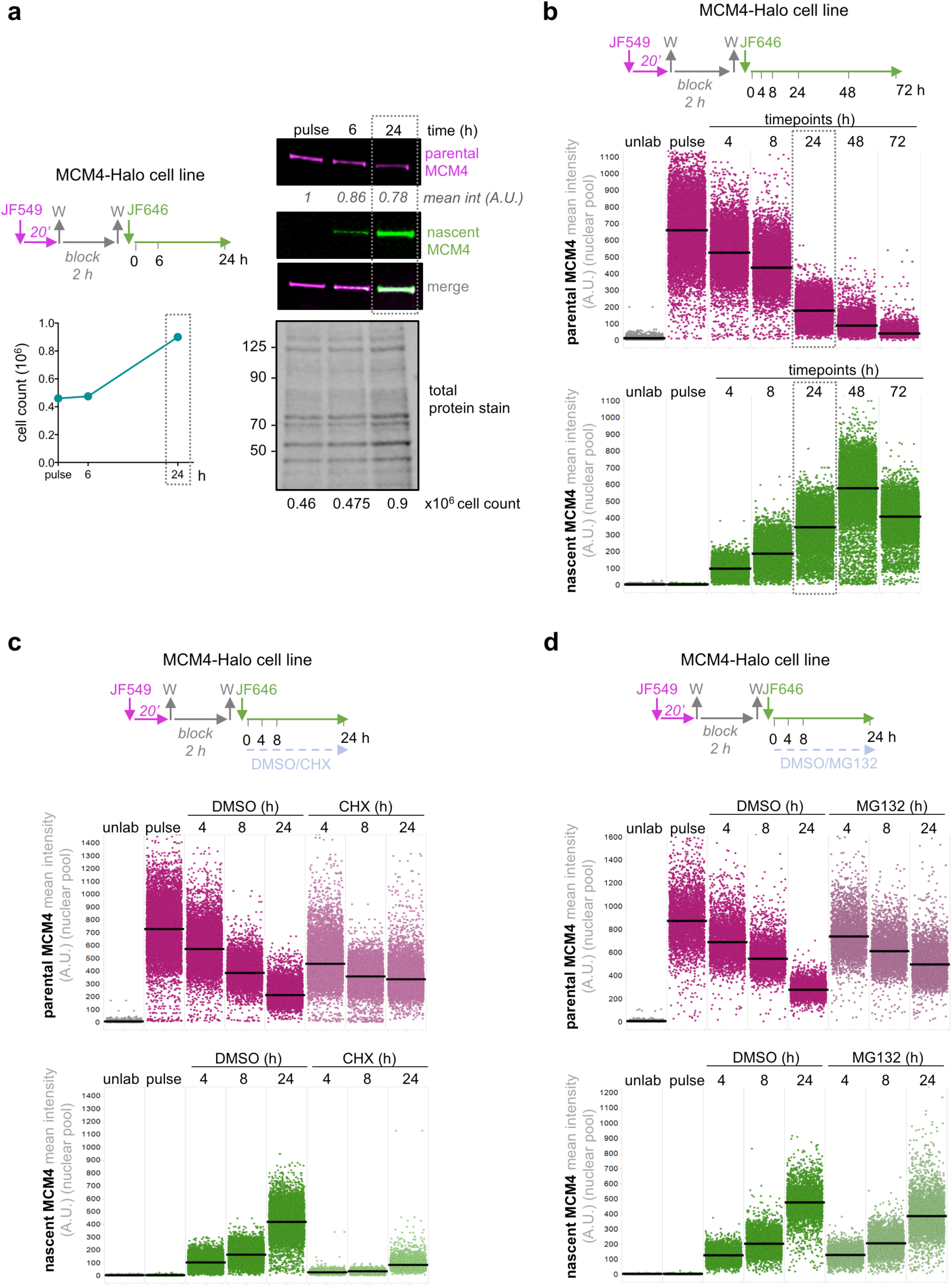
Newly synthesized MCMs constantly replenish the declining pool of recycled parental MCMs. **a,** Left (top), dual-HaloTag labeling protocol in MCM4-Halo cells. Left (bottom), cell count for indicated timepoints. Right, SDS-PAGE of whole cell lysates of MCM4-Halo cells labeled for nascent and parental MCM4 at indicated timepoints (with indicated cell count). Total protein stain as loading control. Box marks the cell doubling time. **b,** Top, dual-HaloTag labeling protocol in MCM4-Halo cells. Middle, QIBC of MCM4-Halo cells immunostained for parental (magenta) MCM4 without pre-extraction before fixation. Bottom, QIBC of MCM4-Halo cells stained for nascent (green) MCM4 without pre-extraction before fixation. Boxes indicate the cell doubling time, horizontal lines are medians. n ≈10000 cells per condition. **c,** Top, dual-HaloTag labeling protocol in MCM4-Halo cells treated as indicated with cycloheximide (CHX; 12.5 μg/ml). Middle, QIBC of MCM4-Halo cells immunostained for parental (magenta) MCM4 without pre-extraction before fixation at indicated timepoints after the indicated treatments. Bottom, QIBC of MCM4-Halo cells stained for nascent (green) MCM4 without pre-extraction before fixation at indicated timepoints after indicated treatments. Horizontal lines are medians. n ≈9000 cells per condition. **d,** Top, dual-HaloTag labeling protocol in MCM4-Halo cells treated as indicated with MG132 (2 μM). Middle, QIBC of MCM4-Halo cells immunostained for parental MCM4 (magenta) without pre-extraction before fixation after indicated treatments. Bottom, QIBC of MCM4-Halo cells stained for nascent MCM4 (green) without pre-extraction before fixation at indicated timepoints after indicated treatments. Horizontal lines are medians. N ≈3500 cells per condition.

**Extended Data Fig. 3.**
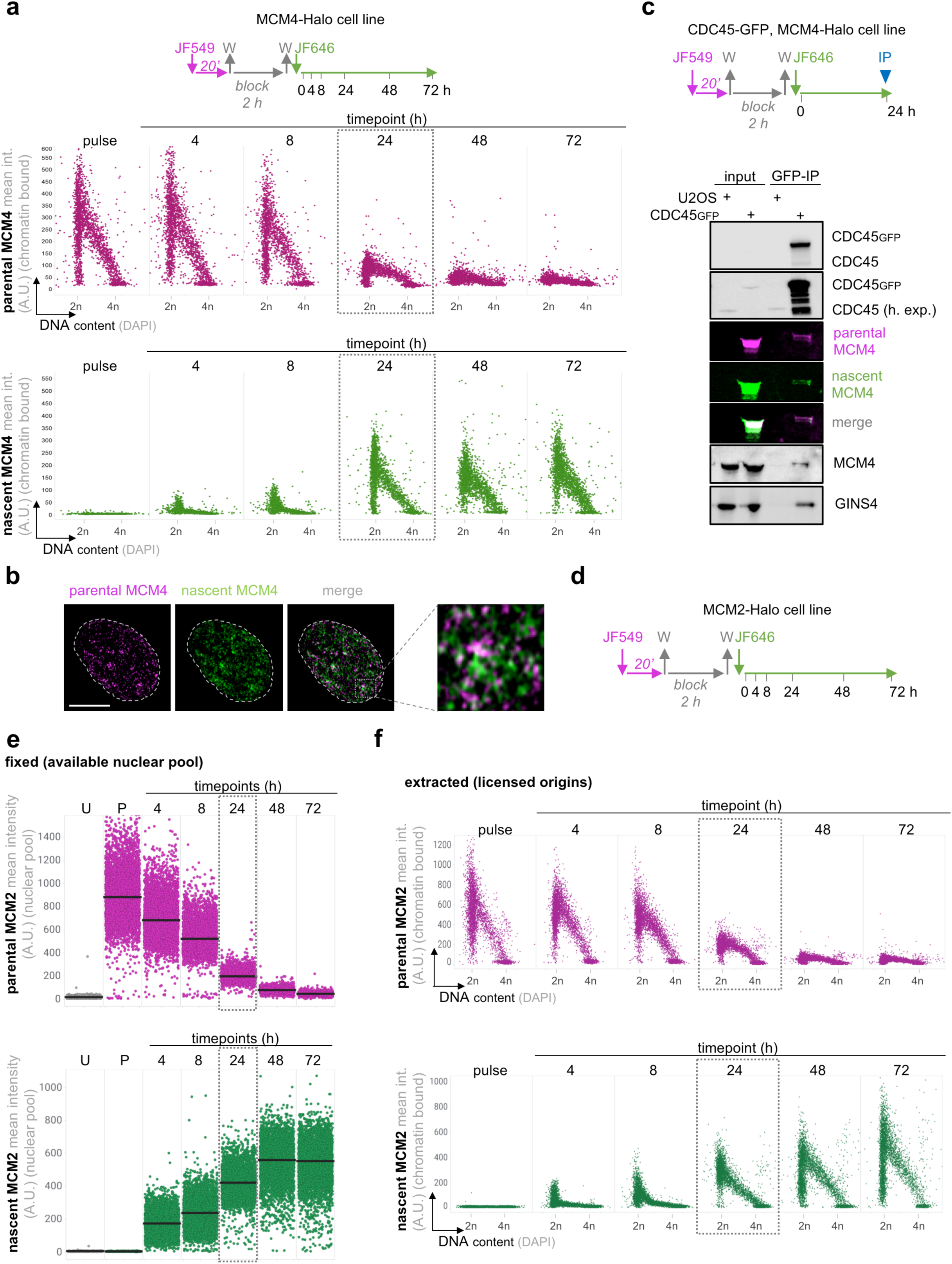
Nascent and parental MCMs are equally proficient in pre-RC licensing and CMG formation. **a,** Top, dual-HaloTag labeling protocol in MCM4-Halo cells. Middle, QIBC of MCM4-Halo cells immunostained for parental MCM4 (magenta) with pre-extraction before fixation. Bottom, QIBC of MCM4-Halo cells stained for nascent MCM4 (green) with pre-extraction before fixation. Boxes mark the cell doubling time. n ≈2000 cells per condition. **b,** Representative confocal images of chromatin-bound parental (magenta) and nascent (green) MCM4 inherited by a daughter cell. Scale bar, 14 μm. **c,** Top, dual-HaloTag labeling protocol in MCM4-Halo cells with endogenously GFP-tagged CDC45. Blue triangle represents collection of lysates for immunoprecipitation (IP). Bottom, GFP-IP of whole cell lysates immunostained before collection for nascent- and parental-MCM4 and then immunoblotted for indicated proteins. **d,** Dual-HaloTag labeling protocol in MCM2-Halo cells. **e,** QIBC of MCM2-Halo cells immunostained for parental MCM2 (magenta) without pre-extraction before fixation. Bottom, QIBC of MCM2-Halo cells immunostained for nascent MCM2 (green) without pre-extraction before fixation. Staining of parental and nascent MCM2 was performed according labeling protocol in (d). Boxes mark the cell doubling time. Horizontal lines are medians. n ≈4400 cells per condition. **f,** Top, QIBC of MCM2-Halo cells immunostained for parental MCM2 (magenta) with pre-extraction before fixation. Bottom, QIBC of MCM2-Halo cells immunostained for nascent MCM2 (green) with pre-extraction before fixation. Box marks the cell doubling time. n ≈3000 cells per condition.

**Extended Data Fig. 4.**
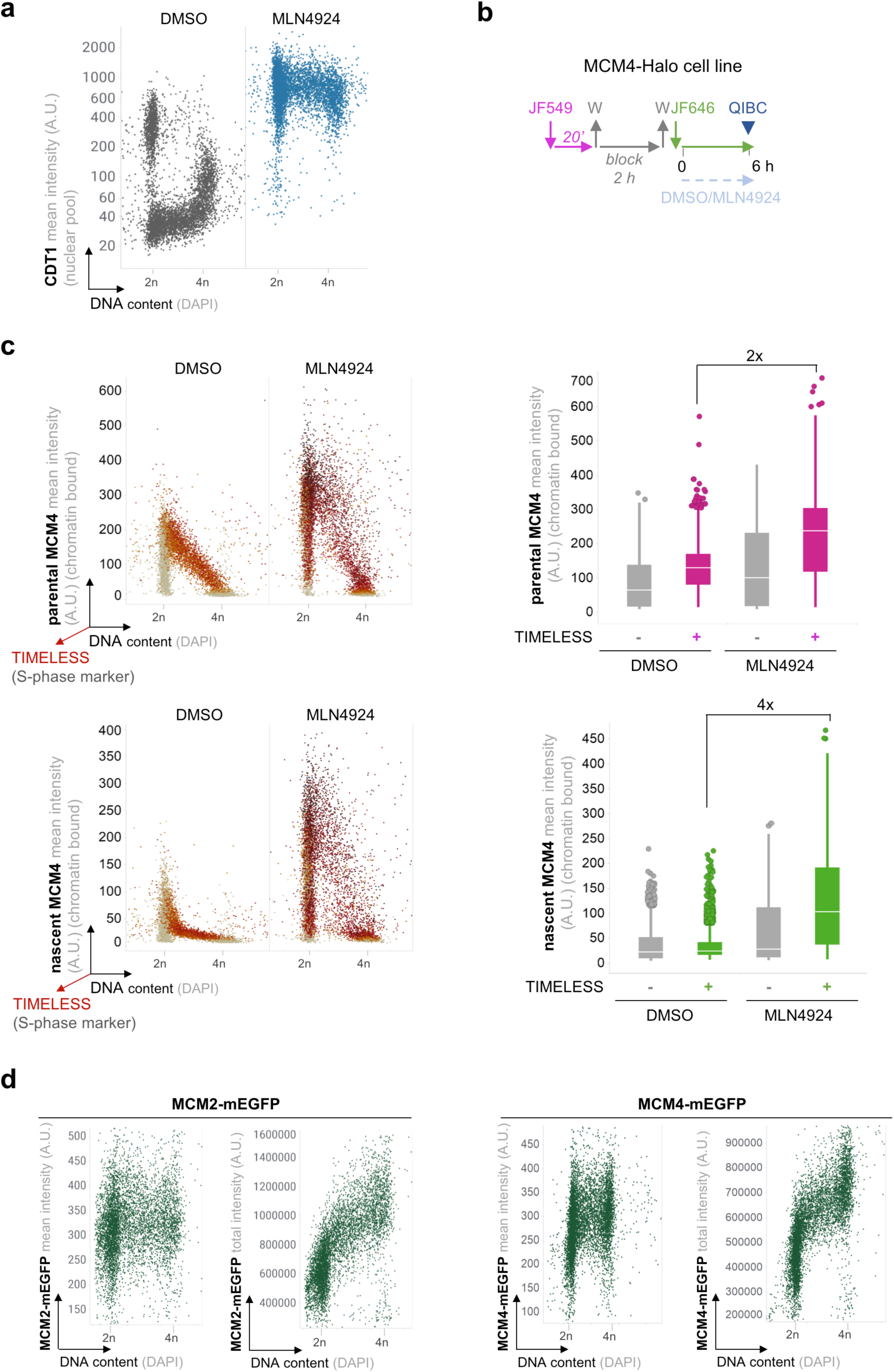
Both nascent and parental MCMs are maintained in licensing-competent mode before cell division. **a,** QIBC of MCM4-Halo cells immunostained for CDT1 and counterstained for DAPI after treatment with DMSO (negative control) or MLN4924 (5 μM; 6 h) without pre-extraction before fixation. n ≈5300 cells per condition. **b,** Dual-HaloTag labeling protocol in MCM4-Halo cells with indicated DMSO or MLN4924 treatments. Blue triangle indicates collection of cells for QIBC. **c,** Top (left), QIBC of MCM4-Halo cells with JF549-labeled parental MCM4 immunostained for TIMELESS (red) and counterstained for DAPI after indicated treatments with pre-extraction before fixation. Top (right), quantification of parental MCM4 fluorescence intensity in TIMELESS-positive or -negative cells after indicated treatments. The center lines in the plots are medians. Bottom (left), QIBC of MCM4-Halo cells with JF646-labeled nascent MCM4, immunostained for TIMELESS and counterstained for DAPI after indicated treatments with pre-extraction before fixation. Bottom (right), quantification of nascent MCM4 in cells TIMELESS-positive or -negative cells after indicated treatments. The center lines in the plots are medians. n ≈7400 cells per condition. **d,** Left, QIBC of cells with endogenously GFP-tagged MCM2 stained with DAPI without pre-extraction before fixation. n ≈8300 cells per condition. Right, QIBC of cells with endogenously GFP-tagged MCM4 stained with DAPI without pre-extraction before fixation. n ≈8300 cells per condition.

**Extended Data Fig. 5.**
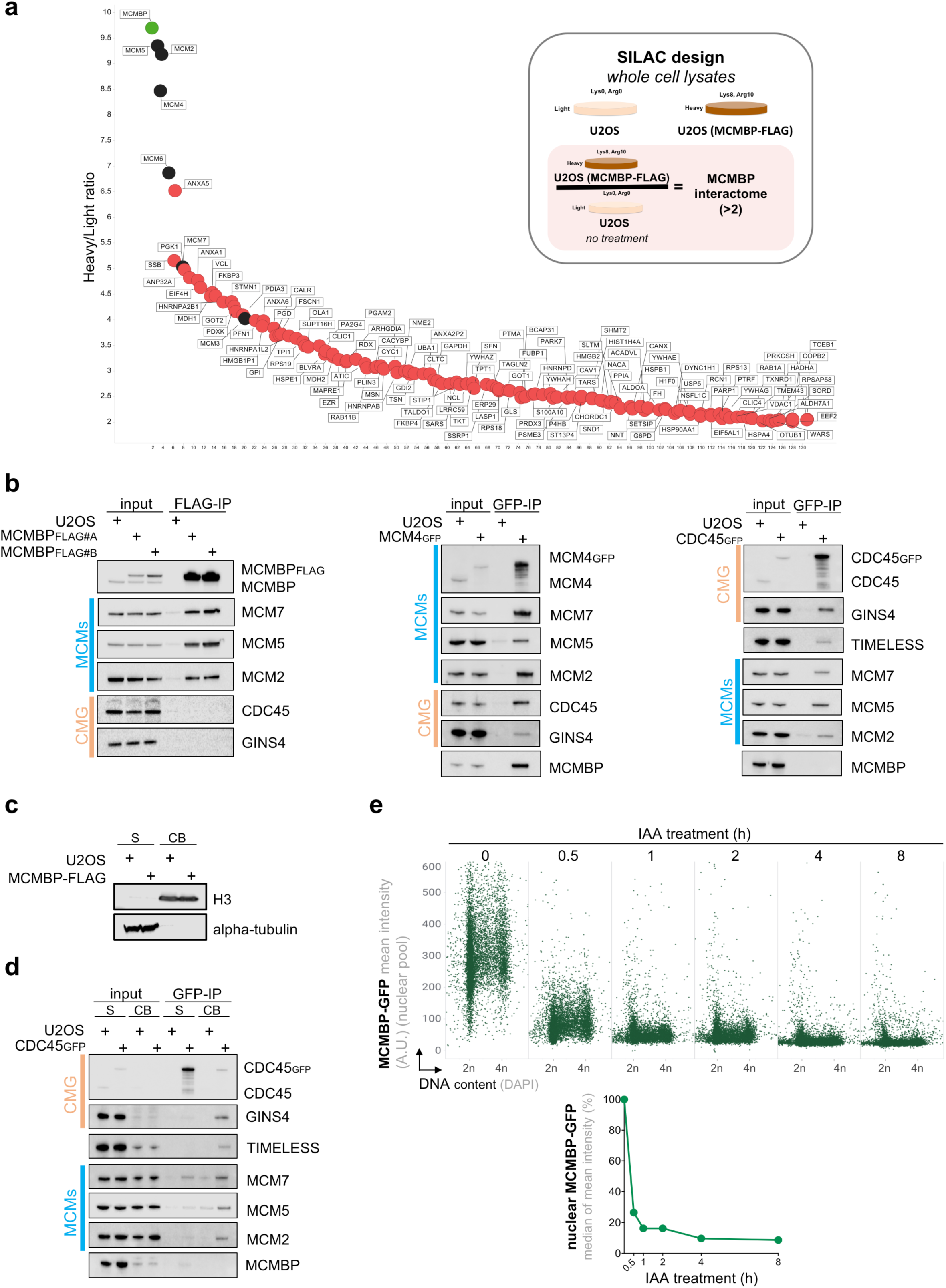
MCMBP associates with CMG helicase independent pool of MCMs. **a,** Interactome of MCMBP obtained upon FLAG-immunoprecipitation (FLAG-IP) from U2OS cells or its derivative ectopically expressing FLAG-tagged MCMBP. FLAG-IP was whole cell extracts (using RIPA lysis buffer with 150 mM NaCl) was analyzed by mass spectrometry. Inset represents SILAC design and criteria for analysis of MCMBP interactome. **b,** Left, FLAG-IP followed by immunoblotting of whole cell extract from U2OS cells or its derivative ectopically expressing FLAG-tagged MCMBP. Middle, GFP-immunoprecipitation (GFP-IP) followed by immunoblotting of whole cell extract from U2OS cells or its derivative endogenously expressing GFP-tagged MCM4. Right, GFP-immunoprecipitation (GFP-IP) followed by immunoblotting of whole cell extract from U2OS cells or its derivative endogenously expressing GFP-tagged CDC45. **c,** Sub-cellular fraction (500 mM NaCl) from U2OS cells or its derivative stably expressing FLAG-tagged MCMBP followed by immunoblotting of H3 or alpha-tubulin. S: supernatant, CB: chromatin-bound. **d,** GFP-IP followed by immunoblotting of sub-cellular fractions (500 mM NaCl) from U2OS cells or its derivative endogenously expressing GFP-tagged MCM4. **e,** Top, QIBC of cells with endogenously GFP-AID-tagged MCMBP stained with DAPI without pre-extraction before fixation at indicated timepoints after auxin (IAA; 0.5 mM) treatment. n ≈6000 cells per condition. Bottom, line plot derived from QIBC results on the top.

**Extended Data Fig. 6.**
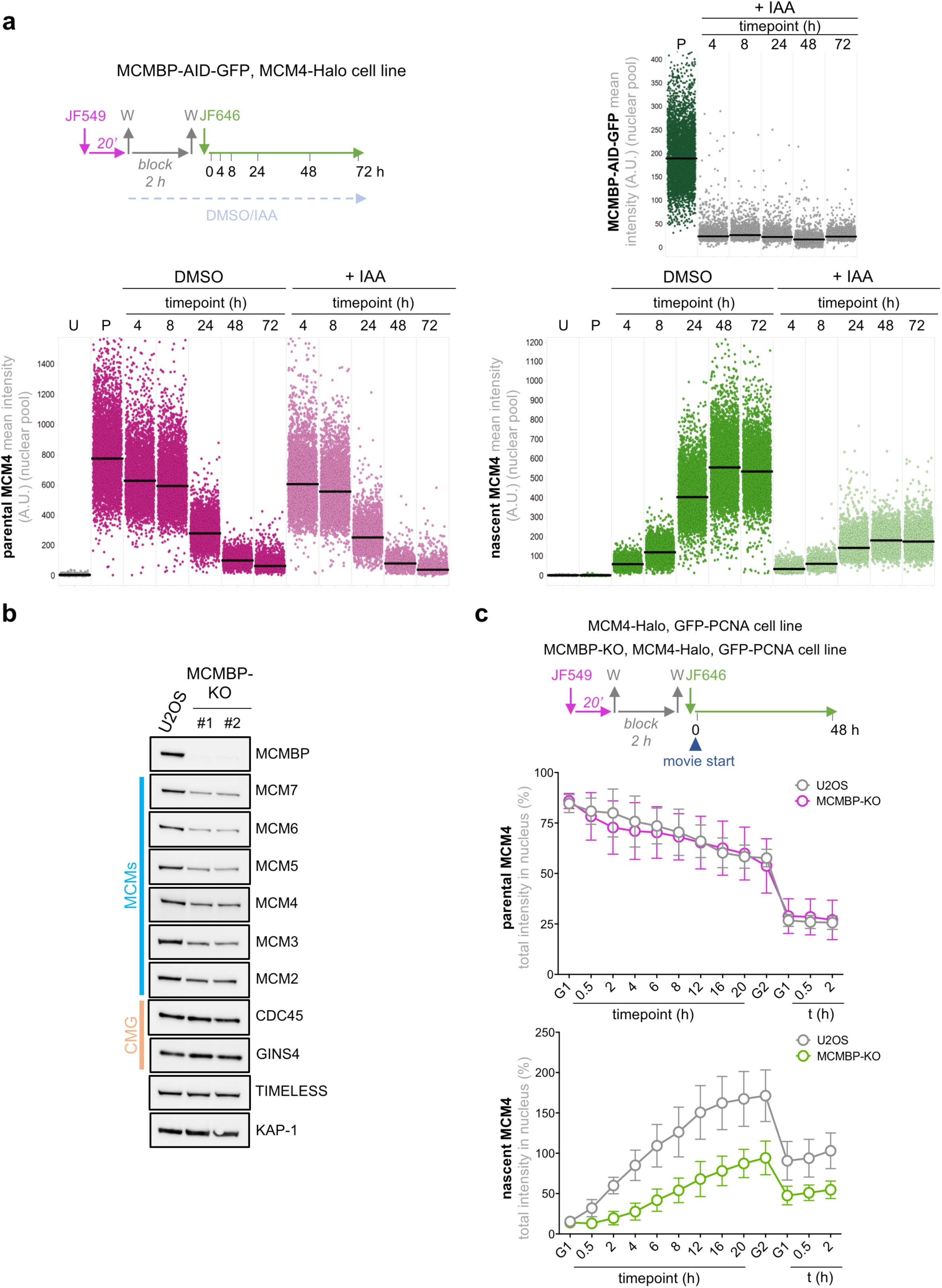
MCMBP fosters nuclear accumulation of nascent but not parental MCMs. **a,** Top (left), dual-HaloTag labeling protocol in MCMBP-AID-GFP cells with endogenously Halo-tagged MCM4 with specified IAA treatment (0.5 mM). Top (right), QIBC of cells MCMBP-AID-GFP cells with endogenously Halo-tagged MCM4 stained with DAPI without pre-extraction before fixation at indicated timepoints after IAA treatment. Bottom (left), QIBC of MCM4-Halo cells stained for parental MCM4 (magenta) without pre-extraction before fixation after indicated treatments. Bottom (right), QIBC of MCM4-Halo cells stained for nascent MCM4 (green) without pre-extraction before fixation after indicated treatments. Horizontal lines medians. U: unlabeled cells, P: pulse. n ≈4000 cells per condition. **b,** Whole-cell extracts from U2OS and MCMBP-KO cells (two independent clones #1 and #2) immunoblotted with indicated antibodies. KAP-1 was used as loading control. **c,** Top, dual-HaloTag labeling protocol as in Fig1c. Total intensities of parental (middle) and nascent (bottom) MCM4 at the start of time-lapse microscopy were considered as 100 percent and the data display relative values for U2OS (MCM4-Halo) and MCMBP-KO (MCM4-Halo) cells. Each data point indicates mean ±SD. n = 15 cells.

**Extended Data Fig. 7.**
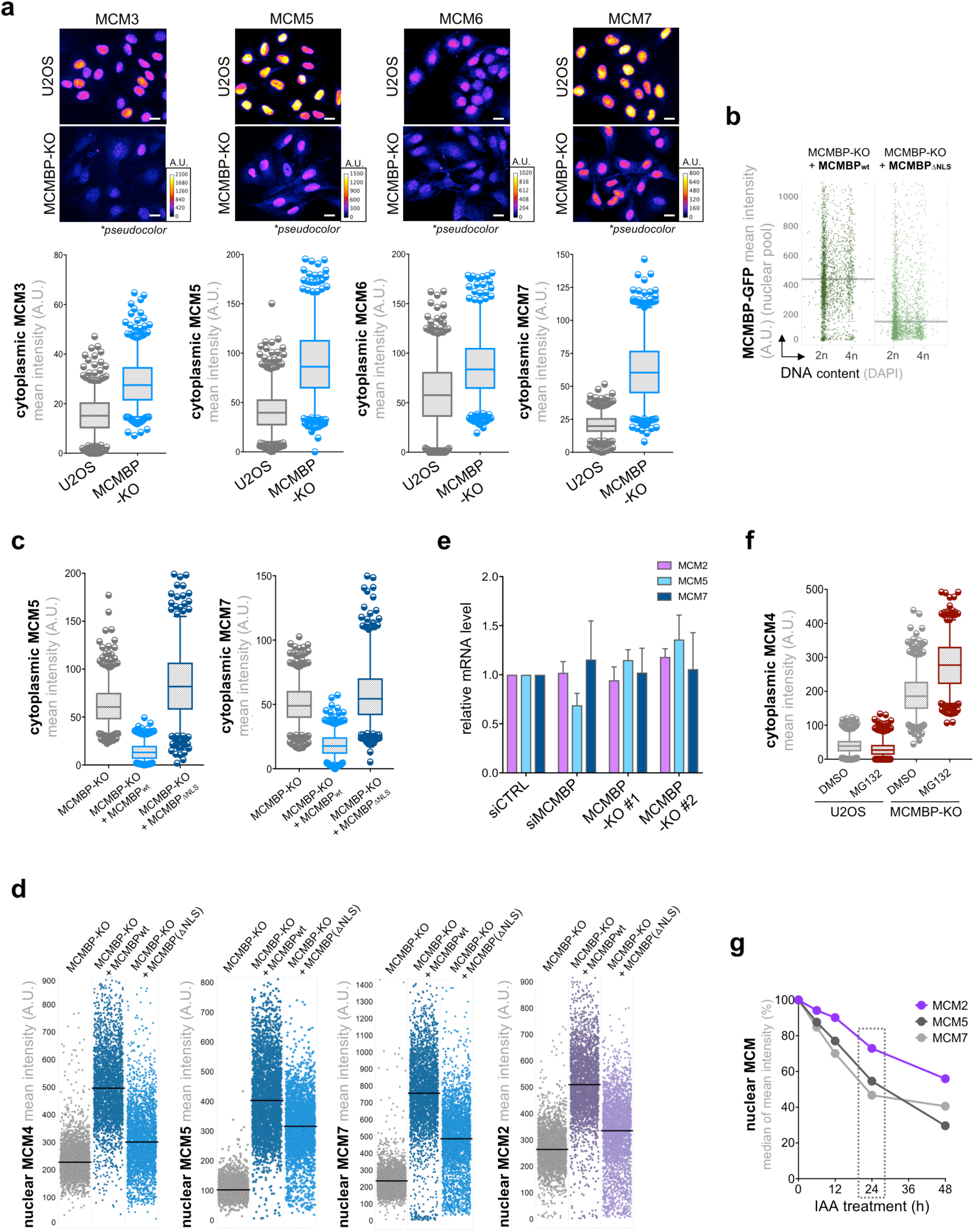
MCMBP possesses an autonomous NLS motif that regulates the rapid nucleo-cytoplasmic shuttling of MCM3-7. **a,** Top, representative images of immunostained MCMs in naïve U2OS and MCMBP-KO cells without pre-extraction before fixation. The color gradient indicates the mean MCM intensity. Scale bar, 20 μm. Bottom, quantification of mean fluorescence intensity (MFI) of cytoplasmic MCMs. The center lines in the plots are medians. The boxes indicate the 25th and 75th centiles, and the whiskers indicate 5 and 95 percent values. **b,** QIBC of MCMBP-KO cells ectopically expressing MCMBP_wt_-GFP or MCMBP_ΔNLS_-GFP stained with DAPI without pre-extraction before fixation. The line represents median. n ≈2700 per condition. **c,** MFI of cytoplasmic MCM5 (left) and MCM7 (right). n = 500 per condition. **d,** QIBC of MCMBP-KO cells or MCMBP-KO cells ectopically expressing MCMBP_wt_-GFP or MCMBP_ΔNLS_-GFP stained for indicated MCMs without pre-extraction before fixation. The center lines in the plots represent the median. n ≈2800 per condition. **e,** RT-PCR analysis of mRNA level for MCM2, MCM5 and MCM7 for indicated cells. mRNA level in control cells was normalized as 100 percent. Data represents mean± SD from 3 technical replicates. **f,** MFI of cytoplasmic MCM4 for indicated cells treated with DMSO or MG132 (2 μM; 6 h) as indicated. **g,** MFI of nuclear MCM2, MCM5 and MCM7 after treating MCMBP-degron cells with IAA (0.5 mM) for indicated timepoints. Each timepoint displays median of mean intensity of nuclear MCM derived from ≈5000 cells. Box marks the cell doubling time.

**Extended Data Fig. 8.**
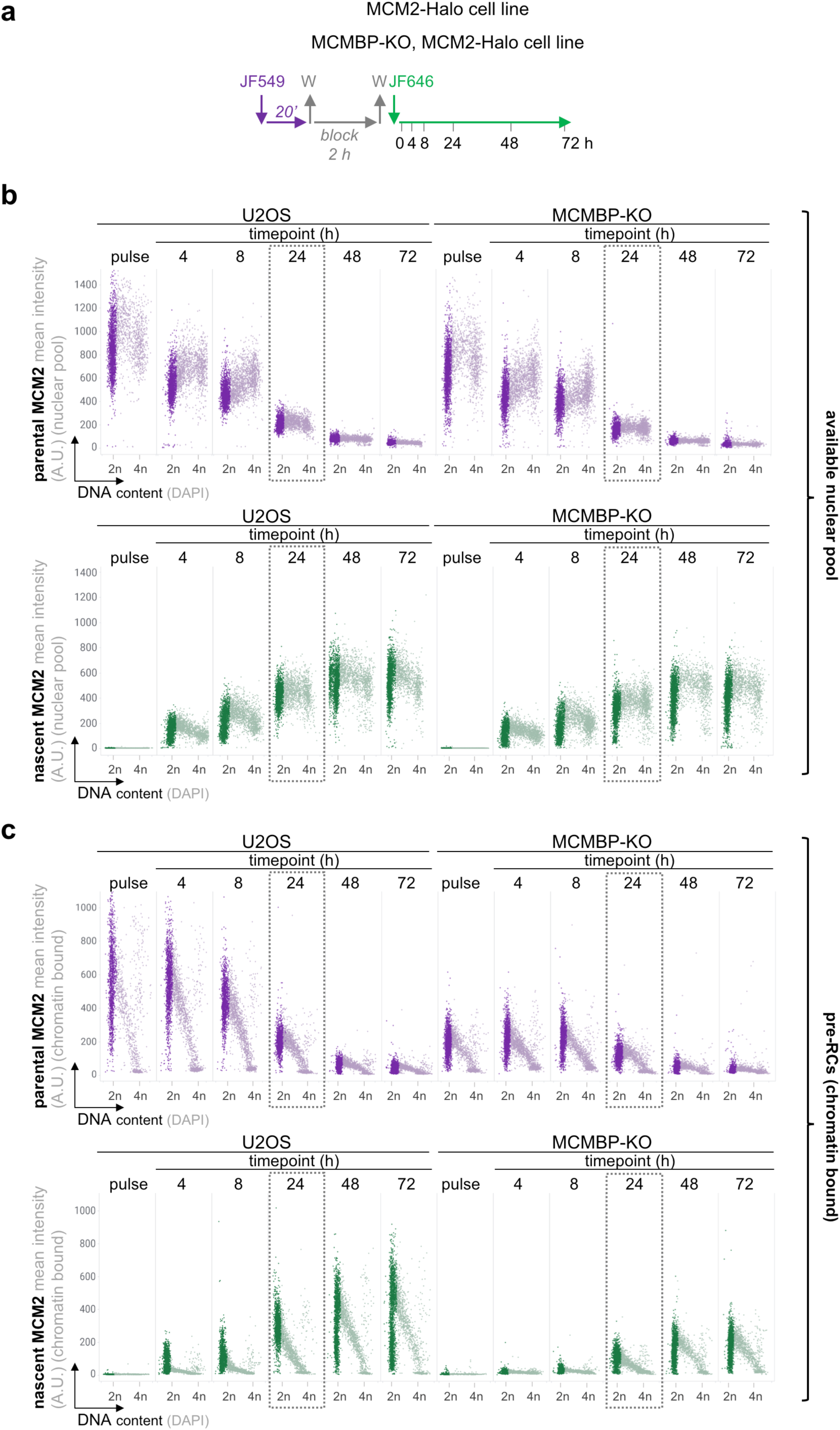
Analysis of the total nuclear and chromatin-bound pool of nascent and parental MCM2 in normal and MCMBP deficient cells. **a,** Dual-HaloTag labeling protocol in U2OS (MCM2-Halo) and MCMBP-KO (MCM2-Halo) cells. **b,** Top, QIBC of U2OS (MCM2-Halo) and MCMBP-KO (MCM2-Halo) immunostained for parental MCM2 (purple) without pre-extraction before fixation for indicated timepoints. Bottom, QIBC of U2OS (MCM2-Halo) and MCMBP-KO (MCM2-Halo) immunostained for nascent MCM2 (green) without pre-extraction before fixation for indicated timepoints. Immunostaining of parental and nascent MCM2 was performed according labeling protocol in a. Boxes mark the cell doubling time and data presented in Fig.3a, b. n ≈3000 cells per condition. **c,** Top, QIBC of U2OS (MCM2-Halo) and MCMBP-KO (MCM2-Halo) immunostained for parental MCM2 (purple) with pre-extraction before fixation for indicated timepoints. Bottom, QIBC of U2OS (MCM2-Halo) and MCMBP-KO (MCM2-Halo) immunostained for nascent MCM2 (green) with pre-extraction before fixation for indicated timepoints. immunostaining of parental and nascent MCM2 was performed according labeling protocol in a. Box marks the cell doubling time and data presented in Fig.3d, e. n ≈2400 cells per condition.

**Extended Data Fig. 9.**
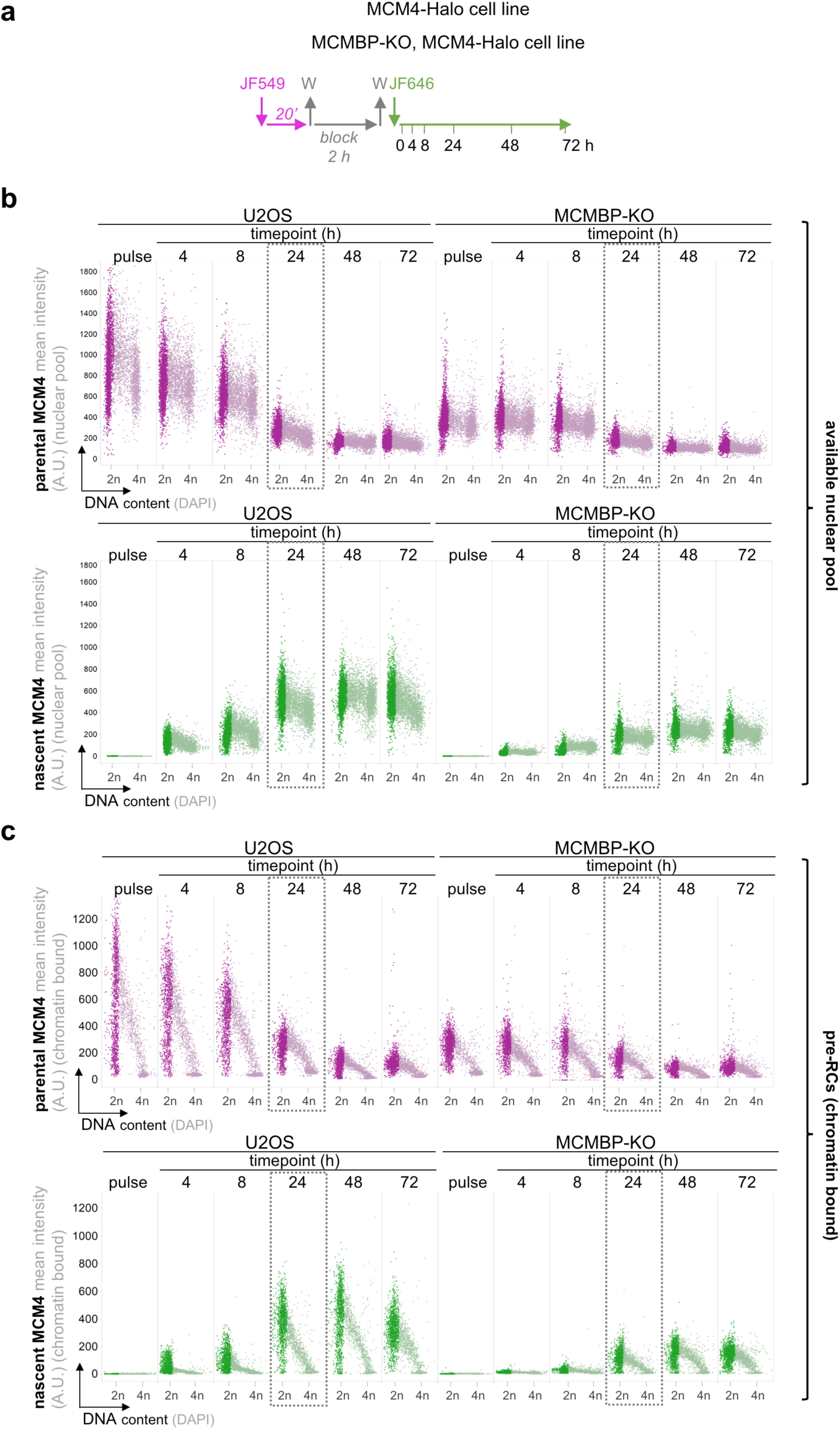
Analysis of the total nuclear and chromatin-bound pool of nascent and parental MCM4 in normal and MCMBP deficient cells. **a,** Dual-HaloTag labeling protocol in U2OS (MCM4-Halo) and MCMBP-KO (MCM4-Halo) cells. **b,** Top, QIBC of U2OS (MCM4-Halo) and MCMBP-KO (MCM4-Halo) immunostained for parental MCM4 (magenta) without pre-extraction before fixation for indicated timepoints. Bottom, QIBC of U2OS (MCM4-Halo) and MCMBP-KO (MCM4-Halo) immunostained for nascent (green) MCM4 without pre-extraction before fixation for indicated timepoints. Staining of parental and nascent MCM4 was performed according labeling protocol in a. Boxes mark the cell doubling time and data presented in Fig.3a, b. n ≈4700 cells per condition. **c,** Top, QIBC of U2OS (MCM4-Halo) and MCMBP-KO (MCM4-Halo) immunostained for parental MCM4 (magenta) with pre-extraction before fixation for indicated timepoints. Bottom, QIBC of U2OS (MCM4-Halo) and MCMBP-KO (MCM4-Halo) immunostained for nascent MCM4 (green) with pre-extraction before fixation for indicated timepoints. Immunostaining of parental and nascent MCM4 was performed according labeling protocol in a. Box marks the cell doubling time and data presented in Fig.3d, e. n ≈2200 cells per condition.

**Extended Data Fig. 10.**
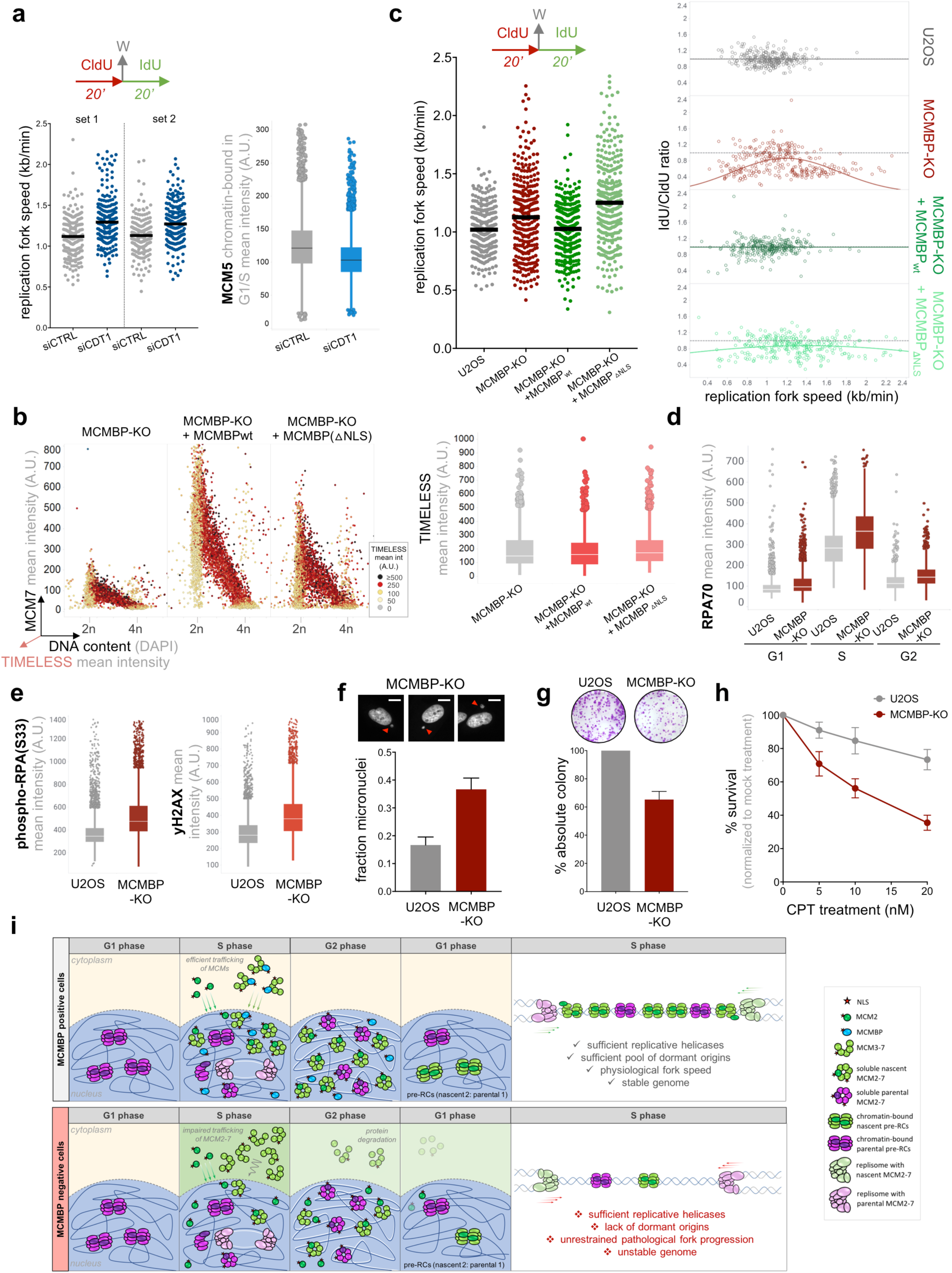
The lack of MCM surplus and paucity of pre-RC licensing in MCMBP deficient cells is associated with increased fork speed and replication stress. **a,** Left (top), DNA fiber labeling protocol. Left (bottom), replication fork speed in cells treated with indicated siRNAs. The center lines in the plots are medians. n = 200 fibers per condition. Right, MFI of chromatin-bound MCM5 in G1/S in cells treated with indicated siRNA. The center lines in the plots are medians. n ≈3800 cells per condition. **b,** Left, QIBC of chromatin-associated MCM7 in indicated cell lines. The color gradient represents the mean intensity of chromatin-bound TIMELESS. n ≈4000 cells per condition. Right, quantification of chromatin-bound TIMELESS. The center lines in the plots are medians. **c,** Left (top), DNA fiber labeling protocol. Left (bottom), replication fork speed in indicated cell lines. The center lines in the plots are medians. n = 300 fibers per condition. Right, individual fork ratio derived from the data in (left) by dividing the length of DNA tracts labeled by IdU and CldU, respectively. The lines represent Gaussian fitting. **d,** QIBC of ssDNA-bound RPA during cell cycle phases in indicated cell lines. The center lines in the plots are medians. n ≈5700 cells per condition. **e,** Left, QIBC of phospho-RPA (S33) in indicated cell lines. Right, QIBC of γH2AX in indicated cell lines. The center line in the plots are medians. n ≈8000 cells per condition. **f,** Frequency of micronuclei formation (500 nuclei per condition) derived from the indicated exponentially growing cell lines and represented as percentage of all counted nuclei per condition. Mean ± SD (from 3 independent biological replicates). **g,** Relative plating efficiency of MCMBP-KO cells compared to naïve U2OS. Mean ± SD, n = 3, technical replicates. **h,** Clonogenic survival of U2OS and MCMBP-KO cells, 10 days after continuous treatment with CPT with indicated concentrations (mean ± SD, n = 3, technical replicates) **i,** A hypothetical model depicting the efficient production, nuclear transport and stable inheritance of MCM2-7, and the role of MCMBP in this process, to ensure optimal levels of origin licensing and replication fork progression in successive cell generations (see text for details).

## References

1 Deegan, T. D. & Diffley, J. F. MCM: one ring to rule them all. Curr Opin Struct Biol 37, 145–151, doi:10.1016/j.sbi.2016.01.014 (2016).

2 Ibarra, A., Schwob, E. & Mendez, J. Excess MCM proteins protect human cells from replicative stress by licensing backup origins of replication. Proc Natl Acad Sci U S A 105, 8956–8961, doi:10.1073/pnas.0803978105 (2008).

3 Alver, R. C., Chadha, G. S. & Blow, J. J. The contribution of dormant origins to genome stability: from cell biology to human genetics. DNA Repair (Amst) 19, 182–189, doi:10.1016/j.dnarep.2014.03.012 (2014).

4 Liang, D. T., Hodson, J. A. & Forsburg, S. L. Reduced dosage of a single fission yeast MCM protein causes genetic instability and S phase delay. J Cell Sci 112 (Pt 4), 559–567 (1999).

5 Orr, S. J. et al. Reducing MCM levels in human primary T cells during the G(0)-->G(1) transition causes genomic instability during the first cell cycle. Oncogene 29, 3803–3814, doi:10.1038/onc.2010.138 (2010).

6 Sakwe, A. M., Nguyen, T., Athanasopoulos, V., Shire, K. & Frappier, L. Identification and characterization of a novel component of the human minichromosome maintenance complex. Mol Cell Biol 27, 3044–3055, doi:10.1128/MCB.02384-06 (2007).

7 Blow, J. J. & Dutta, A. Preventing re-replication of chromosomal DNA. Nat Rev Mol Cell Biol 6, 476–486, doi:10.1038/nrm1663 (2005).

8 Kuipers, M. A. et al. Highly stable loading of Mcm proteins onto chromatin in living cells requires replication to unload. J Cell Biol 192, 29–41, doi:10.1083/jcb.201007111 (2011).

9 Gambus, A. Termination of Eukaryotic Replication Forks. Adv Exp Med Biol 1042, 163–187, doi:10.1007/978-981-10-6955-0_8 (2017).

10 Roseaulin, L. C. et al. Coordinated degradation of replisome components ensures genome stability upon replication stress in the absence of the replication fork protection complex. PLoS Genet 9, e1003213, doi:10.1371/journal.pgen.1003213 (2013).

11 Prasanth, S. G., Mendez, J., Prasanth, K. V. & Stillman, B. Dynamics of pre-replication complex proteins during the cell division cycle. Philos Trans R Soc Lond B Biol Sci 359, 7–16, doi:10.1098/rstb.2003.1360 (2004).

12 Toledo, L. I. et al. ATR prohibits replication catastrophe by preventing global exhaustion of RPA. Cell 155, 1088–1103, doi:10.1016/j.cell.2013.10.043 (2013).

13 Leonhardt, H. et al. Dynamics of DNA replication factories in living cells. J Cell Biol 149, 271–280, doi:10.1083/jcb.149.2.271 (2000).

14 Braun, K. A. & Breeden, L. L. Nascent transcription of MCM2-7 is important for nuclear localization of the minichromosome maintenance complex in G1. Mol Biol Cell 18, 1447–1456, doi:10.1091/mbc.e06-09-0792 (2007).

15 Lin, J. J., Milhollen, M. A., Smith, P. G., Narayanan, U. & Dutta, A. NEDD8-targeting drug MLN4924 elicits DNA rereplication by stabilizing Cdt1 in S phase, triggering checkpoint activation, apoptosis, and senescence in cancer cells. Cancer Res 70, 10310–10320, doi:10.1158/0008-5472.CAN-10-2062 (2010).

16 Taipale, M. et al. A quantitative chaperone interaction network reveals the architecture of cellular protein homeostasis pathways. Cell 158, 434–448, doi:10.1016/j.cell.2014.05.039 (2014).

17 Santosa, V., Martha, S., Hirose, N. & Tanaka, K. The fission yeast minichromosome maintenance (MCM)-binding protein (MCM-BP), Mcb1, regulates MCM function during prereplicative complex formation in DNA replication. J Biol Chem 288, 6864–6880, doi:10.1074/jbc.M112.432393 (2013).

18 Nishiyama, A., Frappier, L. & Mechali, M. MCM-BP regulates unloading of the MCM2-7 helicase in late S phase. Genes Dev 25, 165–175, doi:10.1101/gad.614411 (2011).

19 Kimura, H., Ohtomo, T., Yamaguchi, M., Ishii, A. & Sugimoto, K. Mouse MCM proteins: complex formation and transportation to the nucleus. Genes Cells 1, 977–993 (1996).

20 Ghosh, S., Vassilev, A. P., Zhang, J., Zhao, Y. & DePamphilis, M. L. Assembly of the human origin recognition complex occurs through independent nuclear localization of its components. J Biol Chem 286, 23831–23841, doi:10.1074/jbc.M110.215988 (2011).

21 Scorah, J. & McGowan, C. H. Claspin and Chk1 regulate replication fork stability by different mechanisms. Cell Cycle 8, 1036–1043, doi:10.4161/cc.8.7.8040 (2009).

22 Spies, J. et al. 53BP1 nuclear bodies enforce replication timing at under-replicated DNA to limit heritable DNA damage. Nat Cell Biol 21, 487–497, doi:10.1038/s41556-019-0293-6 (2019).

23 Moreno, A. et al. Unreplicated DNA remaining from unperturbed S phases passes through mitosis for resolution in daughter cells. Proc Natl Acad Sci U S A 113, E5757–5764, doi:10.1073/pnas.1603252113 (2016).

24 Jackson, D. A. & Pombo, A. Replicon clusters are stable units of chromosome structure: evidence that nuclear organization contributes to the efficient activation and propagation of S phase in human cells. J Cell Biol 140, 1285–1295, doi:10.1083/jcb.140.6.1285 (1998).

25 Somyajit, K. et al. Redox-sensitive alteration of replisome architecture safeguards genome integrity. Science 358, 797–802, doi:10.1126/science.aao3172 (2017).

26 Zeman, M. K. & Cimprich, K. A. Causes and consequences of replication stress. Nat Cell Biol 16, 2–9, doi:10.1038/ncb2897 (2014).

27 Jackson, V. In vivo studies on the dynamics of histone-DNA interaction: evidence for nucleosome dissolution during replication and transcription and a low level of dissolution independent of both. Biochemistry 29, 719–731, doi:10.1021/bi00455a019 (1990).

28 Daigh, L. H., Liu, C., Chung, M., Cimprich, K. A. & Meyer, T. Stochastic Endogenous Replication Stress Causes ATR-Triggered Fluctuations in CDK2 Activity that Dynamically Adjust Global DNA Synthesis Rates. Cell Syst 7, 17–27 e13, doi:10.1016/j.cels.2018.05.011 (2018).

29 Matson, J. P. et al. Intrinsic checkpoint deficiency during cell cycle re-entry from quiescence. J Cell Biol 218, 2169–2184, doi:10.1083/jcb.201902143 (2019).

30 Das, M., Singh, S., Pradhan, S. & Narayan, G. MCM Paradox: Abundance of Eukaryotic Replicative Helicases and Genomic Integrity. Mol Biol Int 2014, 574850, doi:10.1155/2014/574850 (2014).

31 Shima, N. et al. A viable allele of Mcm4 causes chromosome instability and mammary adenocarcinomas in mice. Nat Genet 39, 93–98, doi:10.1038/ng1936 (2007).

32 Kunnev, D. et al. DNA damage response and tumorigenesis in Mcm2-deficient mice. Oncogene 29, 3630–3638, doi:10.1038/onc.2010.125 (2010).

## Additional references

33 Koch, B. et al. Generation and validation of homozygous fluorescent knock-in cells using CRISPR-Cas9 genome editing. Nat Protoc 13, 1465–1487, doi:10.1038/nprot.2018.042 (2018).

34 Cong, L. et al. Multiplex genome engineering using CRISPR/Cas systems. Science 339, 819–823, doi:10.1126/science.1231143 (2013).

35 Kosugi, S., Hasebe, M., Tomita, M. & Yanagawa, H. Systematic identification of cell cycle-dependent yeast nucleocytoplasmic shuttling proteins by prediction of composite motifs. Proc Natl Acad Sci U S A 106, 10171–10176, doi:10.1073/pnas.0900604106 (2009).

36 Rappsilber, J., Mann, M. & Ishihama, Y. Protocol for micro-purification, enrichment, pre-fractionation and storage of peptides for proteomics using StageTips. Nat Protoc 2, 1896–1906, doi:10.1038/nprot.2007.261 (2007).

37 Cox, J. et al. Andromeda: a peptide search engine integrated into the MaxQuant environment. J Proteome Res 10, 1794–1805, doi:10.1021/pr101065j (2011).

38 Yamaguchi, K., Inoue, S., Ohara, O. & Nagase, T. Pulse-chase experiment for the analysis of protein stability in cultured mammalian cells by covalent fluorescent labeling of fusion proteins. Methods Mol Biol 577, 121–131, doi:10.1007/978-1-60761-232-2_10 (2009).

39 Ochs, F. et al. 53BP1 fosters fidelity of homology-directed DNA repair. Nat Struct Mol Biol 23, 714–721, doi:10.1038/nsmb.3251 (2016).

40 Rapsomaniki, M. A. et al. easyFRAP: an interactive, easy-to-use tool for qualitative and quantitative analysis of FRAP data. Bioinformatics 28, 1800–1801, doi:10.1093/bioinformatics/bts241 (2012).

